# Distributed processing for action control by prelimbic circuits targeting anterior-posterior dorsal striatal subregions

**DOI:** 10.1101/2021.12.01.469698

**Authors:** Kyuhyun Choi, Eugenio Piasini, Luigim Cifuentes-Vargas, Edgar Díaz-Hernández, Nathan T. Henderson, Manivannan Subramaniyan, Charles R. Gerfen, Marc V. Fuccillo

## Abstract

Fronto-striatal circuits have been extensively implicated in the cognitive control of behavioral output for both social and appetitive rewards. The functional diversity of prefrontal cortical populations is strongly dependent on their synaptic targets, with control of motor output strongly mediated by connectivity to the dorsal striatum. Despite evidence for functional diversity along the anterior-posterior axis of the dorsomedial striatum (DMS), it is unclear how distinct fronto- striatal sub-circuits support neural computations essential for action selection. Here we identify prefrontal populations targeting distinct DMS subregions and characterize their functional roles. We first performed neural circuit tracing to reveal segregated prefrontal populations defined by anterior/posterior dorsomedial striatal target. We then probed the functional relevance of these parallel circuits via *in vivo* calcium imaging and temporally precise causal manipulations during a feedback-based 2-alternative choice task. Single-photon imaging revealed circuit-specific representations of task-relevant information with prelimbic neurons targeting anterior DMS (PL::A- DMS) uniquely encoded choices and responses to negative outcomes, while prelimbic neurons targeting posterior DMS (PL::P-DMS) encoded internal representations of value and positive outcomes contingent on prior choice. Consistent with this distributed coding, optogenetic inhibition of PL::A-DMS circuits strongly impacted choice monitoring and behavioral control in response to negative outcomes while perturbation of PL::P-DMS signals impaired task engagement and strategies following positive outcomes. Di-synaptic retrograde tracing uncovered differences in afferent connectivity that may underlie these pathways functional divergence. Together our data uncover novel PL populations engaged in distributed processing for action control.

**SUMMARY:**

- Prelimbic cortex engages A- and P-DMS via distinct circuits
- PL::A-DMS and PL::P-DMS pathways encode divergent aspects of a simple goal-directed task
- PL::A-DMS pathways shape responding to negative outcomes via multiple mechanisms
- PL::P-DMS pathways guide engagement and choices in response to positive outcomes
- Afferent connectomes of PL neurons defined by A-P DMS target are distinct

## INTRODUCTION

Value-based decision-making requires a complex series of neural computations - the integration of success and failure, the proper attribution of actions to temporally displaced outcomes and the monitoring of context and underlying task structure. One hypothesis posits that inputs for this decision-making process are represented across forebrain excitatory populations, with their integration in the striatum serving as an early step in action selection^1^. Consistent with a topographical organization of afferent inputs^2–4^, striatum exhibits functional segregation along its anatomical axes, with the dorsoventral direction segregating reward and motor processes and medial-lateral domains supporting goal-sensitive and habitual action strategies^5^. However, substantially less work has considered striatal function along the anterior-posterior (A-P) axis^6–10^ despite early retrograde studies pointing to a unique longitudinal (A-P) organization of cortical- striatal inputs^11^.

Seminal studies in rat provided the first evidence of functional segregation along the striatal A-P axis, with posterior dorsomedial striatum (P-DMS) lesions disrupting both the initial acquisition and post-training execution of instrumental conditioning, in particular modulation in responding according to action-outcome association^8, 9^. In contrast, the importance of the anterior dorsomedial striatum (A-DMS) in goal-directed choice remained uncertain, with opposing results for pharmacological inactivation and excitotoxic lesions^8, 9, 12^. Optogenetic manipulations of specific spiny projection neuron subtypes within the A-DMS have implicated this subregion in supporting flexible responses during reversal learning^13^, consistent with pharmacological manipulations of anterior caudate in marmosets^14^. In contrast, the anterior dorsolateral striatum (DLS) supports a protein synthesis-dependent consolidation of newly learned actions^15^. Finally, a growing body of evidence has implicated the rodent striatal tail, the most caudal subregion, in behavioral responses to aversive stimuli and psychostimulants^16–18^.

The prefrontal cortex exerts cognitive control over mammalian behavior via extensive afferent integration and widespread downstream connectivity^19^. Analysis of prefrontal populations accounting for downstream synaptic targets has revealed pathway-specific functional differences for prefrontal control of social-spatial rewards^20^, reward anticipation^21^, and choice directions^22^. The prelimbic region of the prefrontal cortex has been hypothesized to support goal-directed action by encoding short-term memories necessary for subsequent action-outcome associations in dorsal striatum^23^. Specific targeting of prelimbic-striatal pathways has extended this view, demonstrating persistent neural coding of value essential for choice behavior^24^ and the mediation of response inhibition during tasks requiring sustained attention^25^. Finally, DREADD-mediated inhibition of PL neurons projecting to either anterior or posterior striatal subregions has uncovered involvement in instrumental learning^6, 7, 10^.

Here we systematically explore the function of PL pathways projecting along the A-P striatal axis via integration of mono- and di-synaptic viral circuit tracing, single neuron calcium imaging, statistical modeling of neural coding properties, and target-specific optogenetic manipulations. Retrograde tracing from A/P-DMS subregions revealed non-overlapping PL populations which exhibited unique encoding of behavioral variables over multiple time scales essential for shaping efficient action selection and execution. Target- and temporally- specific optogenetic manipulations confirmed the functional divergence of these fronto-striatal pathways, with PL::A- DMS pathways supporting choice monitoring and responding to negative outcomes and PL::P- DMS pathways supporting engagement and responding to positive outcomes. Together, our results provide novel insight into the distributed nature of fronto-striatal pathways for decision making.

## RESULTS

### Anatomical architecture of fronto-striatal pathways along the anterior-posterior striatal axis

To characterize prefrontal cortex connectivity along the anterior-posterior striatal axis, we injected a mix of AAV5-CamKII::GFP-Cre and AAVdj-EF1a::Flex-Synaptophysin-mRuby virus into prelimbic cortex (Fig. 1a), confirming that synaptic inputs from PL were widely spread along the full anterior-posterior extent of DMS (Fig. 1b). To address whether these widespread projections arose from *en passant* connectivity or distinct PL afferents, we utilized two orthogonal retrograde circuit tracers, with EnvA G-deleted rabies virus EGFP injected in A-DMS, and Alexa647- conjugated Cholera toxin subunit-B (CTB) injected in P-DMS (Fig. 1c). This design minimized fiber of passage contamination of PL::P-DMS pathways while traversing A-DMS. Using CTIP2 immunostaining as a guide, we found cell bodies of both retrogradely labeled populations largely in prelimbic layers II/III and more sparsely in layers V/VI ^26^ (Fig. 1f-i). Regardless of layer, these populations were distinct (2.2*±*0.5% overlap) and spatially separated, forming a characteristic sub-layer structure with PL::A-DMS populations localized to superficial layer II/III and PL::P-DMS populations found in deeper layer II/III (Fig. 1e). These results were replicated using spectrally distinct CTB tracers (Fig. S1a-c), confirming the existence of distinct PL cortical populations defined by A/P-DMS subregions and revealing a similar anatomical organization for most striatal afferents originating in other brain regions (Fig. S1d-j).

**Figure 1.**
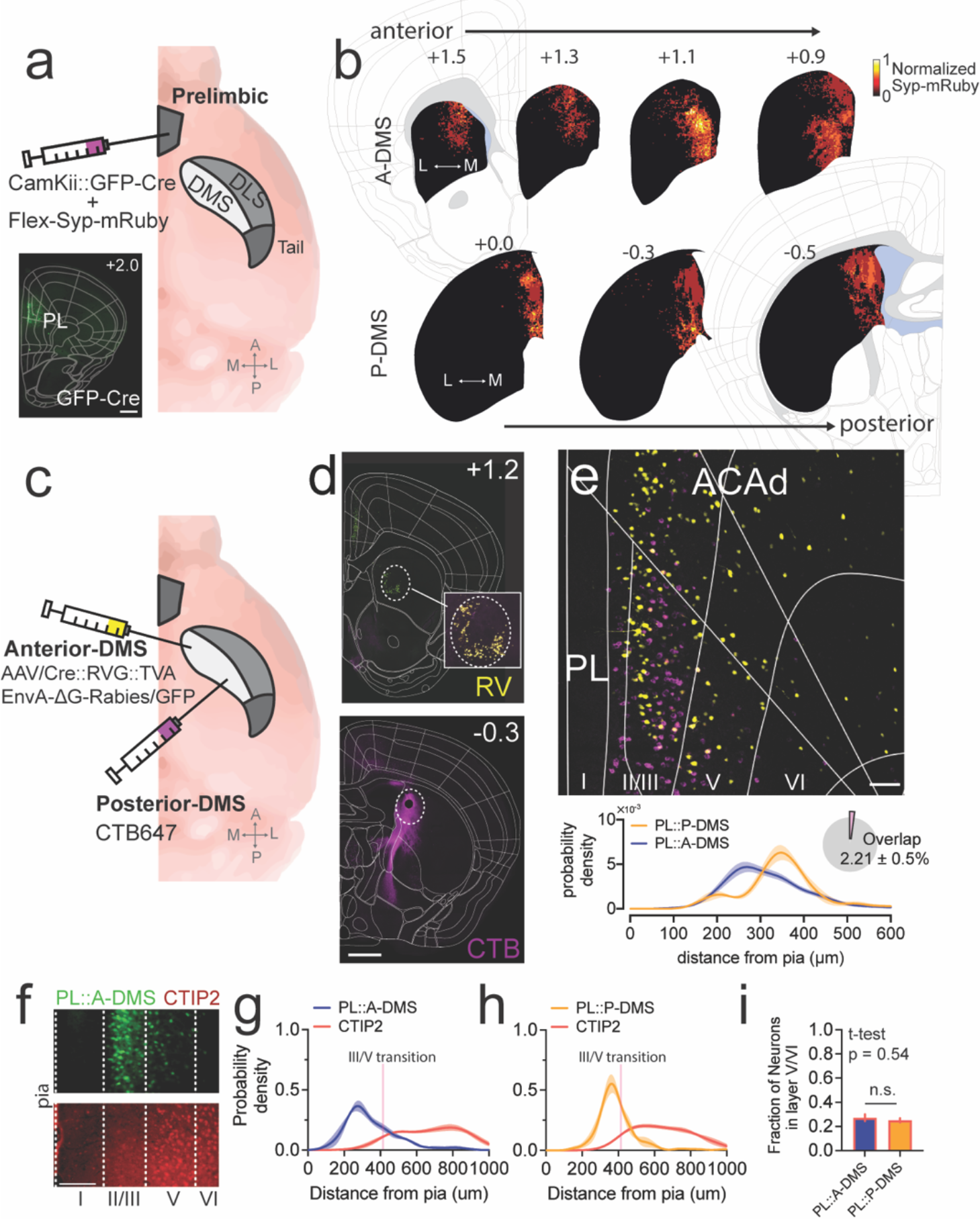
Distinct PL neuron populations defined by anterior/posterior dorsomedial striatal (DMS) target. a) Approach for anterograde tracing of PL-DMS excitatory projections with synaptic terminal marker Synaptophysin-mRuby (inset shows GFP-Cre expression at PL target site. b) Superimposed striatum images (top: A-DMS, bottom: P-DMS, left to right increasingly posterior) showing averaged fluorescent intensity of Synaptophysin-mRuby inputs from PL along anterior-posterior axis (n= 4). c) Schematic demonstrating dual-color retrograde tracing strategy using trans-synaptic rabies virus (A-DMS) and Alexa647-conjugated CTB (P-DMS). d) Coronal sections showing injection sites (top: A-DMS, Bottom: P-DMS). scale bar, 500 μm. Number in upper left corner indicates A/P coordinate from bregma. e) Representative image of dorsomedial prefrontal cortex (top) and quantification (kernel density estimate) of neuronal density from the pia (bottom), with relative proportion of overlapping double-labeled neurons (inset). scale bar, 100 μm (n= 4). ACAd, dorsal part of anterior cingulate cortex. f) Example image showing prelimbic area from EnvA-ΔG-rabies virus tracing of A-DMS (top) co-stained with CTIP2 (bottom). Scale bar 100 μm.g) Quantification of neuronal density from pia of PL::A-DMS and CTIP2+ populations (n= 3). h) Quantification of neuronal density from pia of PL::P-DMS and CTIP2+ populations (n= 3). Pink line in g,h represents average layer III-V transition as visualized by compact CTIP2 staining. i) Fraction of GFP labeled neurons located in compact CTIP2+ deep cortical layers.

### Assessing neural activity in PL::DMS pathways during a goal-directed choice task

This unique circuit architecture could serve to either carry similar neural signals to distinct striatal regions or alternatively support divergent neural processing for the control of action selection. To explore these possibilities, we investigated neural coding of task-relevant information within PL::A-DMS and PL::P-DMS populations during a goal-directed choice task. Mice were trained on a 3-poke chamber where the center port initiated a choice period, requiring a lateral left/right decision. In any given trial, choosing a predetermined side led to the delivery of a reward with 85% chance and no outcome otherwise, while choosing the opposite port led to punishment tone with 85% chance and no outcome otherwise (Fig. 2d). The identity of the rewarded side (or “contingency”) was changed whenever mice made 8 correct choices over the latest 10 trials, to assess flexible responding. As previously reported, mice choices were based on previous outcome feedback, with a strong influence of prior trial on current choice (Fig. S2b)^27^. We performed 1-photon (1-p) single neuron calcium (Ca^2+^) imaging of retrogradely-labeled PL neurons expressing GCaMP7f during this task. Given the minimal fiber-of-passage overlap with standard retrograde tracers (Fig. S1a-c), we injected retroAAV2-EF1a::3XFLAG-Cre into either A-DMS or P-DMS, together with AAV1-hSyn::FLEX-jGcamp7f into PL to gain access to both PL populations in separate animals (Fig. 2a-c). Using this approach, we recorded Ca^2+^ activity of 465 PL::A-DMS neurons and 586 PL::P-DMS neurons.

**Figure 2.**
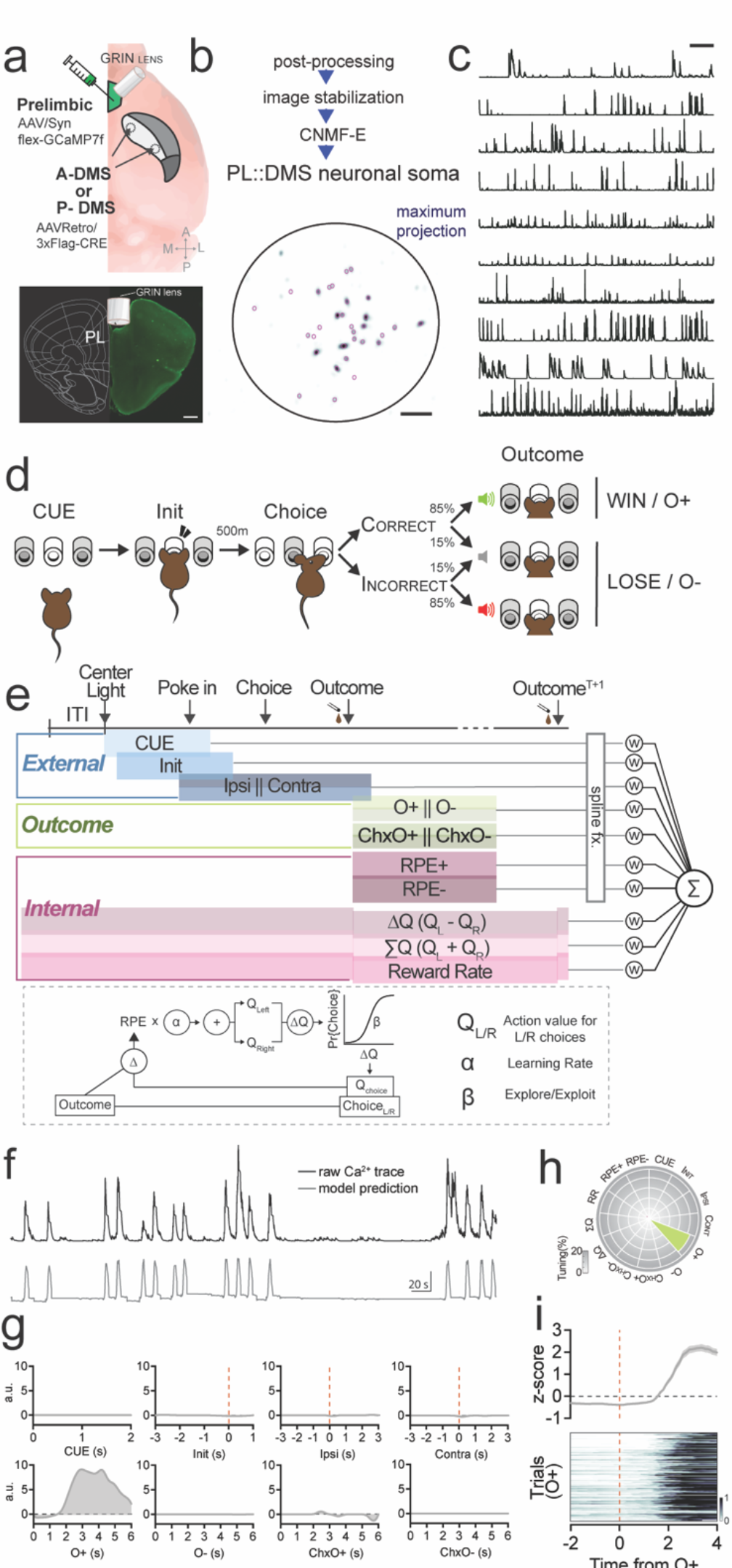
Quantifying neural coding of a value-based operant task. a) Schematic showing viral injection strategy to label pathway specific PL neurons for 1-photon calcium imaging (top) and representative image for GRIN lens location (bottom). scale bar, 500μm. b) MIN1PIPE workflow for extraction of calcium signal from identified ROIs (bottom). scale bar, 50 pixels. c) Representative raw Ca^2+^ traces from 10 PL::A-DMS neurons. Scale bar, 1min. d) Schematic of trial structure showing mice initiating trials via sustained (500 ms) center port entry, followed by left/right choice within 3 sec. Subsequent reward is delivered from center port. e) Schematic drawing of design matrix for neural encoding model showing behavioral predictors for sensorimotor components, outcomes and internal representations of value (top, see table 1). Reinforcement learning model for estimating internal value predictors *(bottom).* f) Example raw Ca^2+^ trace (*top, black*) and output of encoding model (*bottom, gray*). g) Temporally expanded predictors (kernels) from example neuron exhibiting strong O+ modulation. h) Tuning plot of same neuron. i) Peri-event time histogram (PETH, *top*) and trial-by-trial neuronal activity (*bottom*) aligned by O+.

**Table 1.** Details on model predictors. Window extent PRE/POST indicates the extent of the kernel window before/after the event to which the predictor is tethered. N of splines is the size of the spline basis used for the predictor. The basis functions are cubic B- splines. The spline knots are placed at regular intervals within the kernel window; additionally, four knots are placed at each end of the window to enable flexibility in the value of the kernel and all its derivatives at the endpoints. The degree of the splines and the number of the knots determines the number of elements in the basis spline set, reported in the fourth column.]

| Predictor | Window extent PRE (s) | Window extent POST (s) | Number of splines |
| --- | --- | --- | --- |
| start cue (CUE) | 0 | 2 | 12 |
| Self-initiation (Init) | 3 | 1 | 12 |
| Choice LEFT (Lf) | 3 | 3 | 20 |
| Choice RIGHT (Rt) | 3 | 3 | 20 |
| Outcome NEGATIVE (O-) | 0 | 6 | 20 |
| Outcome POSITIVE (O+) | 0 | 6 | 20 |
| Choice RIGHT x Outcome NEGATIVE (Rt x O-) | 0 | 6 | 20 |
| Choice RIGHT x Outcome POSITIVE (Rt x O+) | 0 | 6 | 20 |
| Reward prediction error (RPE), positive (RPE+) | 0 | 6 | 20 |
| Reward prediction error (RPE), negative (RPE-) | 0 | 6 | 20 |
| Reward rate (RR) | (continuous predictor) |  |  |
| $Q(\text{LEFT}) + Q(\text{RIGHT}) (\Sigma Q)$, | (continuous predictor) | | |
| $Q(\text{LEFT}) - Q(\text{RIGHT}) (\Delta Q)$ | (continuous predictor) | | |

To analyze neural activity, we designed a linear encoding model based upon task-relevant regression predictors capturing actions, sensory input, resulting outcomes and model-based estimations of internal value state (Fig. 2e). External sensorimotor variables included trial start cue (CUE), self-initiation (Init), and Ipsilateral/Contralateral (Ipsi/Cont) choice. Outcomes were divided into positive (O+) and negative (O-), as well as interactions of these terms with prior choice, a potential neural signal for credit assignment (Ch x O+, Ch x O-). Local reward rate over the last 5 trials was included as a proxy for task engagement. Finally, we estimated internal value representations with a standard Q-learning model, which proved strongly predictive of future animal choice in our experiments (Fig. S2a,c)^27, 28^. Latent variables inferred with the Q-learning scheme were included in the neural encoding model as predictors representing trial-by-trial choice values (*Δ*Q, *Σ*Q) and reward prediction errors (RPE+/-). Regression parameters were fit via elastic-net penalized maximum likelihood (Fig. 2e-i; see Methods for details on model design and fitting).

We applied this encoding model to both PL::A-DMS and PL::P-DMS Ca^2+^ imaging data, measuring total model fit quality by calculating the fraction of Ca^2+^ signal variance explained (FVE). At a cut-off threshold of 5% FVE, our model fit ∼39% of total PL::A-DMS neurons and ∼30% of total PL::P-DMS neurons (Fig. 3a-b). To quantify neuronal tuning to specific behavioral variables, partial models lacking the related predictors were fit to Ca^2+^ data. The difference in FVE between the full and the partial model defined a tuning index for the given variables. For an initial overview, we grouped predictors into external (CUE, Init, Ipsi, Contra), internal (*Δ*Q, *Σ*Q, RPE+, RPE-, RR) and outcome (O+, O-, Ch x O+, Ch x O-) categories (Fig. 2e)., discovering that the PL::A-DMS pathway was biased towards representation of external variables, the PL::P-DMS pathway was biased towards representation of internal values, and both pathways shared encoding of task outcome (Fig. 3c-d).

**Figure 3.**
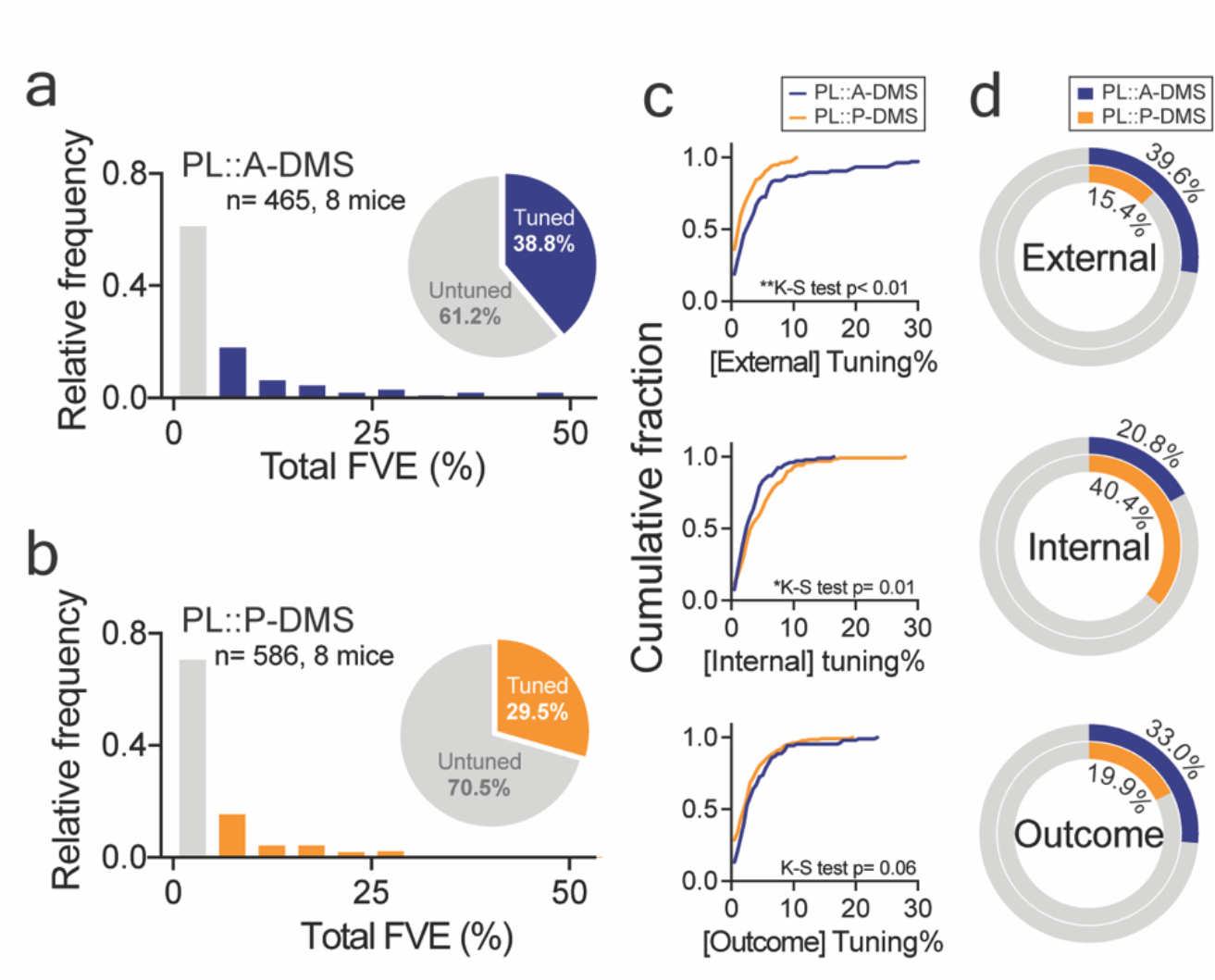
Divergence of neural coding for PL pathways defined by A/P DMS target. a,b) Histogram of binned total FVE distribution for all neurons from (a) PL::A-DMS or (b) PL::P-DMS. Grey bars denote non-task tuned population (<5% FVE); colored bars (blue, PL::A-DMS; orange, PL::P-DMS) denote task-tuned neurons. Pie charts showing the proportion of task tuned neurons for both PL::DMS pathways (insets). c) Cumulative distribution of tuning indices for grouped behavioral variables (see text): external (top), internal (middle), outcome (bottom). Plots are restricted to task-tuned neurons. d) Proportion of highly tuned neurons (>5% FVE) for each behavioral category from each pathway, external (top), internal (middle), outcome (bottom).

### PL::A-DMS and PL::P-DMS neural populations encode distinct and complementary components of value-based behavior

We initially focused on the external bias of PL::A-DMS and asked whether neuronal tuning was specific to sensory or motor events preceding action selection. By breaking down our encoding analysis to the level of individual predictors, we found that the majority of external event modulation in the PL::A-DMS pathway was driven by choice-tuned neurons (Fig. S3a), although PL::A-DMS carried more information than PL::P-DMS not only for choice (Fig. 4d,e, Fig. S3n-r) but also trial start cue (center port light; Fig. S3d-h) and initiation (Fig. S3i-m). We found that PL::A-DMS encoded both ipsilateral and contralateral choices (Fig.4d, Fig. S3b, q).

**Figure 4.**
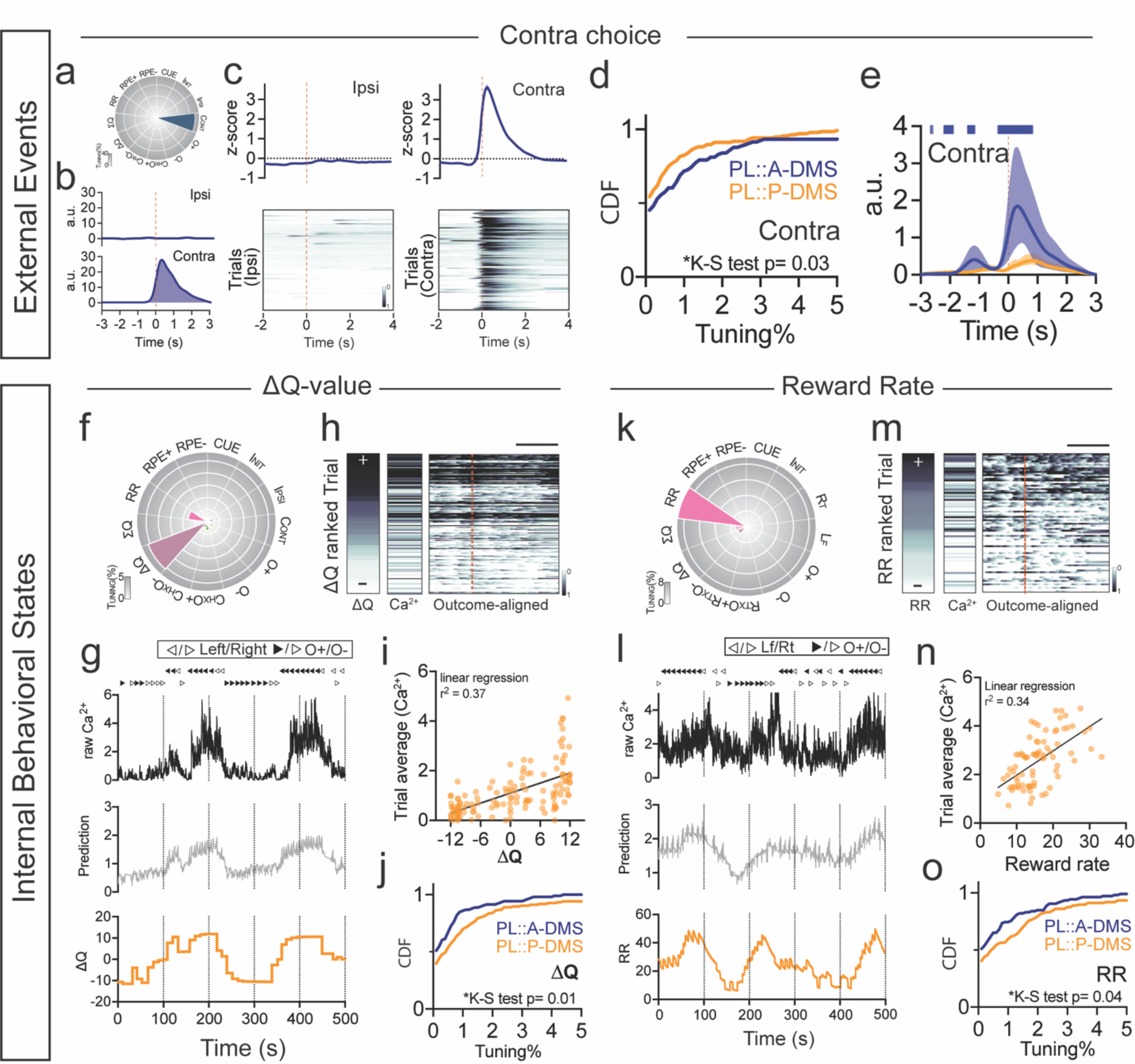
Preferential encoding of choice in PL::A-DMS, internal value signals in PL::P- DMS. a) Tuning plot showing representative contralateral choice tuned neuron from PL::A-DMS. b) Representative kernels corresponding to ipsi (*top*) and contra (*bottom*) choices. c) z-scored PETH (*top*) and trial-by-trial neuronal activity (*bottom*) from Ipsi (*left*)/Contra (*right*) choice. d) Cumulative distribution of contra choice tuned neurons in both PL-DMS pathways. e) Averaged contralateral choice kernels for both PL-DMS pathways. Solid line denotes root-mean-squared; shaded area denotes 95% confidence interval. Colored-bar on top indicates significant mean- displacement on each timepoint between pathways. f) Tuning plot for representative ΔQ tuned neuron from PL::P-DMS. g) Raw Ca^2+^ trace (black, *top*), model prediction (gray, *middle*) and trial- by-trial ΔQ (orange, *bottom*). Choices and outcomes at top (direction of triangle, left/right choice; filled/blanked, O+/O-). h) Trial-by-trial transient Ca^2+^ signals (middle, trial average; right, outcome aligned) ranked by ΔQ (left). scale bar, 2 sec. i) Scatter plot showing linear correlation between ΔQ and trial average of Ca^2+^ waveform. j) Cumulative distribution for ΔQ tuned neurons in both PL-DMS pathways. k) Tuning plot showing representative RR tuned neuron from PL::P-DMS. l) Raw Ca^2+^ trace (black, *top*), model prediction (gray, *middle*) and RR (5-prior trial average, orange, *bottom*) with choice/outcome information at top. m) Trial-by-trial transient Ca^2+^ signals (middle, averaged; right, outcome aligned) ranked by RR (left, averaged). scale bar, 2 sec. n) Scatter plot showing linear correlation between local RR and trial average of Ca^2+^ transients. o) Cumulative distribution for RR tuned neurons in both PL-DMS pathways.

Our tuning index is a compact measure of the degree to which task-related variables are represented in neural activity. Nonetheless, it only captures the overall coding strength and is not sensitive to the precise temporal evolution of neural responses, which could present interesting differences between pathways regardless of their relative tuning level. To address this, we analyzed the event-associated kernels inferred by our encoding model, which estimate the average calcium activity transient elicited by specific behavioral events, after accounting for overlapping transients from other event types. Analysis of choice kernels revealed that PL::A- DMS neurons exhibited robust phasic activity starting around choice execution, although the magnitude of this modulation was on average stronger for choices contralateral to the recording site (Fig. 4e, Fig. S3c, r).

Next, we investigated target-specific PL differences for the representation of internal values, a key driver of decision-making in the absence of task-relevant sensory information. Our reinforcement learning model of choice behavior provides trial-by-trial estimates of the difference between choice values (ΔQ, ipsi vs. contra), the sum of choice values (*Σ*Q) as well as positive and negative reward prediction errors (+/-RPEs). Besides these metrics, we included in our encoding model a local reward rate over the last five trials (RR) to capture the strength of engagement in this self-initiated task. We found that the PL::P-DMS pathway more strongly encoded these internal value estimates (Fig. 3c,d middle), with the strongest drivers being neurons modulated by the difference in action values (ΔQ; Fig. 4j) and those whose activity strongly tracked with the local reward rate (RR; Fig. 4o). Interestingly, our encoding model robustly captured the slow shifting baseline of PL::P-DMS calcium activity that in a subset of neurons scaled with increasing Q-value difference or local reward rate (Fig. 4g,h,l,m) despite lacking clear event-related modulation (Fig. S4e,f). One exception to this dominance of PL::P- DMS for value-related information was for negative RPEs, for which PL::A-DMS pathways demonstrated strong modulation of outcome signals by violated reward expectation (Fig. S4a). Overall however, these data imply that PL::P-DMS pathways more strongly represent temporally integrated internal measures of value than PL::A-DMS pathways.

Finally, we examined how these distinct PL pathways responded during behavioral outcomes, uncovering three general patterns. First, we noted a brief (∼1s) response immediately following all positive outcomes that was similar in calcium waveform between PL pathways but found in a greater proportion of PL::A-DMS neurons (Fig. S5a-h). We also observed neural activity modulated by the interaction of positive outcome and prior choice (Ch x O+; Fig. 5a-c). Interestingly, we found that outcome-related neural signals that were contingent on prior choice were better represented in PL::P-DMS than in PL::A-DMS populations (Fig. 5d). Second, the temporal kinetics of these interaction-associated signals were distinct between pathways, with Ch x O+ signals in PL::P-DMS pathways persisting for several seconds beyond outcome, as compared to briefer Ch x O+ signals in PL::A-DMS neurons (Fig. 5e). Third, we observed robust neuronal responses to negative outcomes that were almost exclusively encoded by the PL::A- DMS neurons (Fig. 5f-i). These signals exhibited a slow and persistent increase following the absence of reward, which occurred at contingency switches, random unrewarded trials or during brief exploratory choice periods (Fig. 5j). Together, these data reveal a distributed representation of outcomes by PL::DMS pathways, with prolonged activation of PL::A-DMS neurons encoding negative outcomes and PL::P-DMS neurons encoding positive outcomes contingent on prior choice.

**Figure 5.**
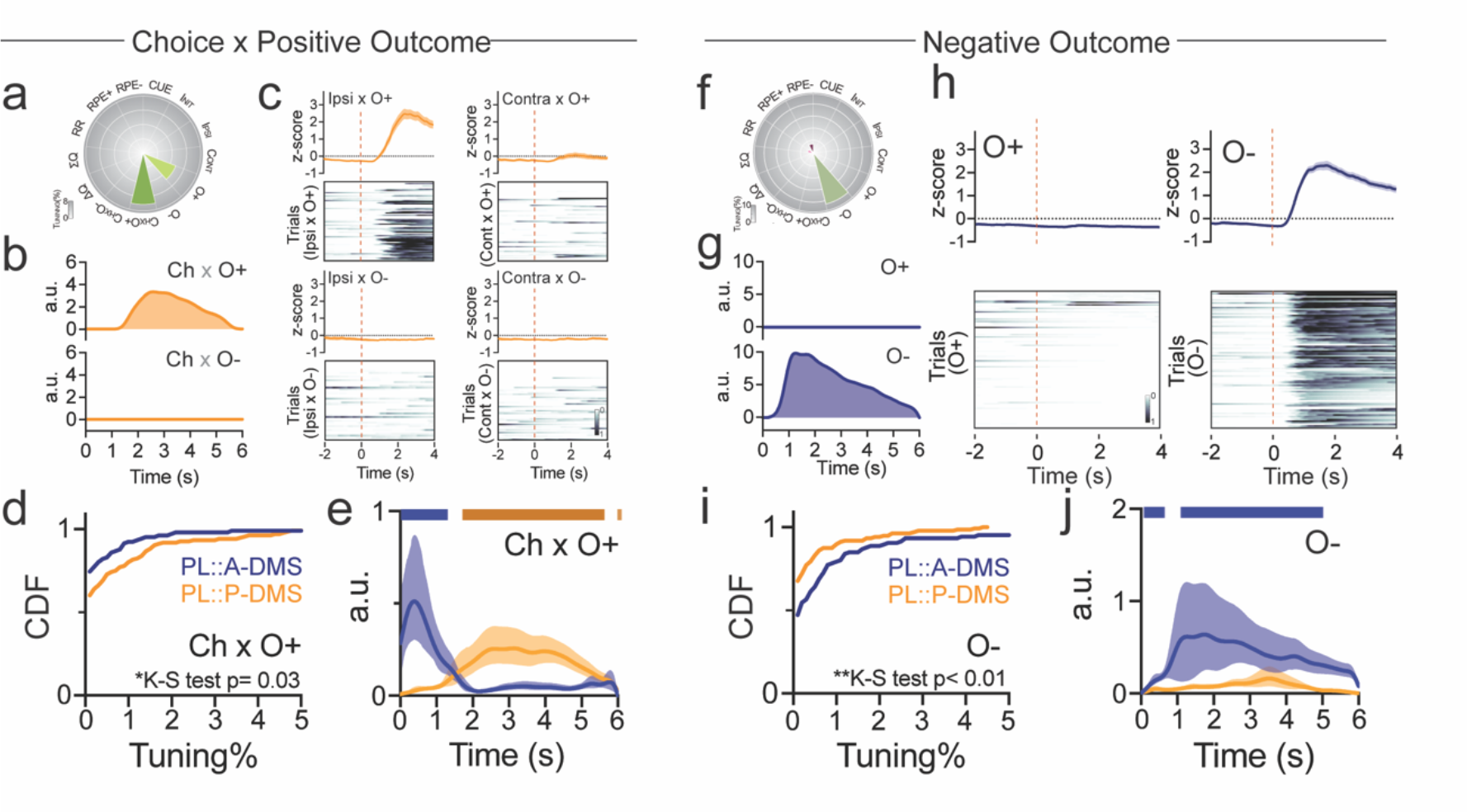
Divergent encoding of outcome by PL::DMS pathways. a) Representative tuning plot of Ch x O+ tuned neuron from PL::P-DMS. b) Inferred kernels corresponding to Ch x O+ (*top*), Ch x O- (*bottom*). c) z-scored PETH (*top* in each panel) and trial-by-trial neuronal activity (*bottom* in each panel) to 4 possible choice outcomes, aligned to outcome delivery. d) Cumulative distribution for Ch x O+ tuning from task-tuned neurons of both PL::DMS pathways. e) Averaged Ch x O+ kernels show pathway-distinct temporal properties. Solid line denotes root-mean-squared; shaded area denotes 95% confidence interval. Colored-bar on top indicates significant mean-displacement on each timepoint between pathways. f) Representative tuning plot for O- tuned neuron from PL::A-DMS. g) Inferred kernels corresponding to O+ (*top*), O- (*bottom*). h) z-scored PETH (*top*) and trial-by-trial neuronal activity (*bottom*) corresponding to types of outcomes (*left*, O+; *right*, O-). i) Cumulative distribution for O- tuning from task-tuned neurons of both PL::DMS pathways. j) Averaged O- kernels for both PL::DMS pathways.

Thus far, our data highlight a unique fronto-striatal architecture defined by A-P striatal target that encodes complementary aspects of relevant external and internal behavioral parameters observed during our value-based task. Our neural coding analysis makes several predictions about pathway-specific behavioral functions: 1. PL::A-DMS choice activity may shape current choice selection/execution or instead provide an action-monitoring signal; 2. PL::P-DMS neurons encode temporally integrated signals for local reward rate and action value that may drive task engagement; 3. the persistent choice x positive outcome activity in PL::P-DMS could be used to drive positive reinforcement behavior; 4. PL::A-DMS negative outcome modulated neurons could be used to implement choice strategies following negative outcome.

### PL::A-DMS pathways mediate future choice valuation, but not current choice execution

To evaluate whether these divergent patterns of neural coding resulted in distinct functional contributions, we performed striatal subregion-specific optogenetic inhibition of PL terminals. We bilaterally injected PL cortex with AAV5-CamKII::NpHR3.0-EYFP, or AAV5-hSyn::EGFP for controls, and implanted 200µm fiber optics bilaterally in either the A-DMS or P-DMS (target sites in Fig. S6a, b). We designed two distinct light delivery protocols to assess the contribution of these fronto-striatal circuits during choice and at outcome. We predicted that PL::A-DMS choice activity might either have a role in the selection/execution of current actions or instead provide an efference copy of the selected action that could be linked to resulting outcomes, thereby influencing future action selection. We also predicted that manipulation of PL::P-DMS pathways would have no effects on choice selection or motor performance, consistent with their lack of choice modulation. To test these predictions, we activated NpHR from initiation through choice on a random 30% subset of trials (Fig. 6, Fig.S7). To analyze effects on choice selection, we took advantage of the strong dependence on prior trial outcomes^27, 28^, analyzing win-stay and lose-stay probabilities (see Methods). We found no evidence that optogenetic inhibition of PL::A-DMS throughout the choice period had any impact upon ongoing action selection (Fig. S7a, Fig S8a,b). To analyze effects on motor performance, we examined choice latency (the time from center port initiation to choice selection), observing no effect of optogenetic inhibition on latency distributions (Fig. S7b). We next analyzed the influence of choice-associated optogenetic suppression on subsequent action selection and performance, finding increased lose-stay behavior following choice activity suppression in prior trials for PL::A-DMS pathways (Fig. 6a, Fig S8g,h). No subsequent trial effect was found for motor performance (Fig. 6b; cf. Fig S8e, k for GFP control). Consistent with our population coding data, optogenetic inhibition of PL terminals in P-DMS had no effect on either choice selection or execution for current or subsequent trials (Fig. 6c, d; Fig. S7c, d; Fig S8c, d, I, j). Overall, these causal manipulations complement the neural coding analysis, suggesting that choice-epoch activity in PL::A-DMS is not related to action planning or execution, but instead provides an efference copy of actions for subsequent valuation.

**Figure 6.**
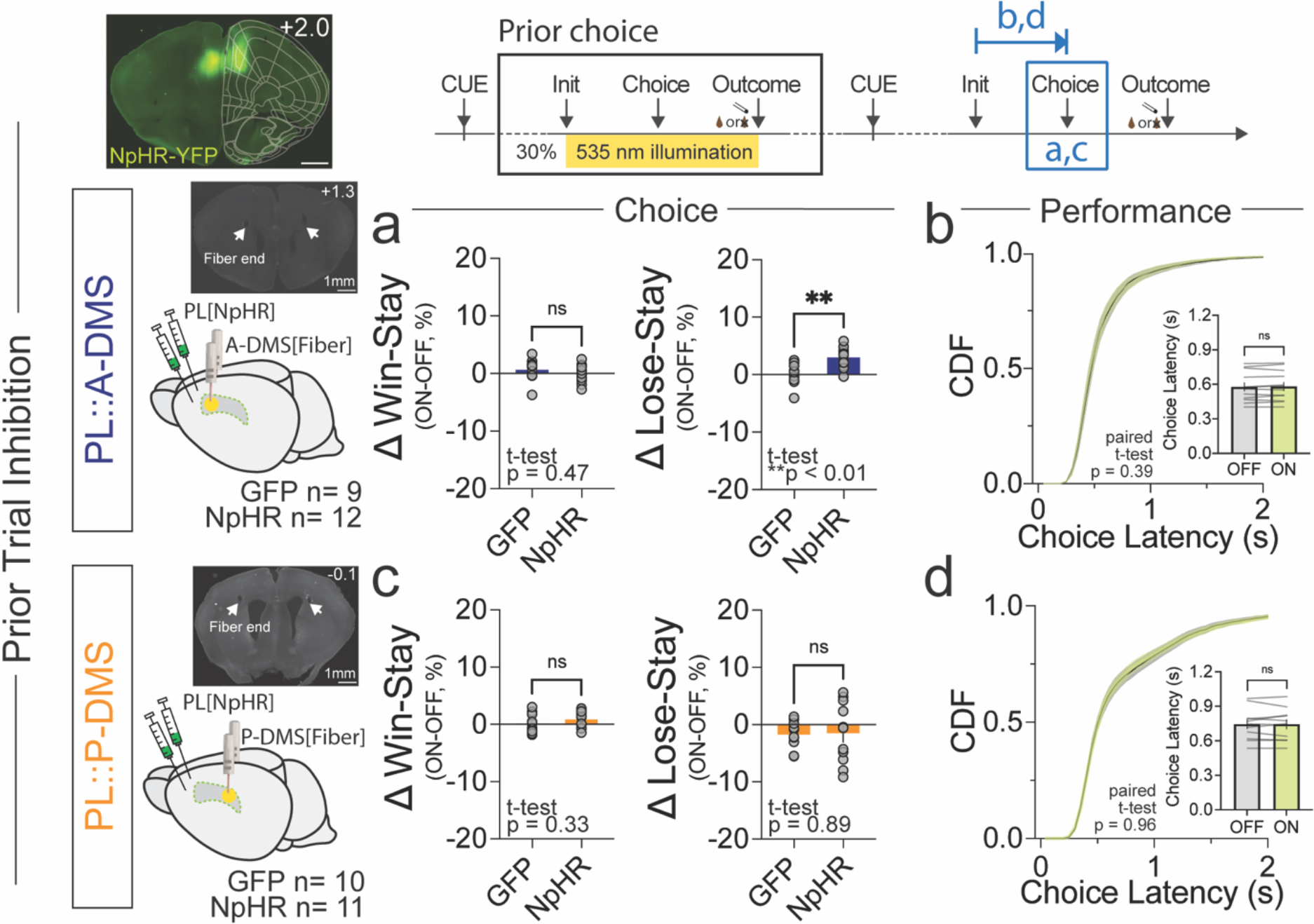
Optogenetic suppression of choice-associated PL::A-DMS activity impairs subsequent choice selection without impacting current trials. (*top, left*) Representative section of PL injection site. (*top, right*) schematic of optogenetic manipulation (yellow bar) with respect to behavioral measures (blue). a) Comparison of ΔWin-Stay (*left*) and ΔLose-Stay (*right*) between NpHR and GFP (control) groups, when light was delivered in the previous choice epoch to PL terminals in A-DMS. b) Cumulative distribution of choice latencies and average of choice latency (inset) following light ON versus OFF trials from NpHR inhibition of PL::A-DMS. c) Comparison of ΔWin-Stay (left) and ΔLose-Stay (right) between NpHR and GFP (control) groups, when light was delivered in the previous choice epoch to PL terminals in P-DMS. d) Cumulative distribution of choice latencies and average of choice latency (inset) following light ON versus OFF trials from NpHR inhibition of PL::P-DMS.

### Temporally integrated PL::P-DMS neural activity supports task engagement

PL::P-DMS pathways were found to strongly encode action value differences and local reward rates, two temporally integrated measures of recent task outcome. As the slow dynamics of these neural signals precluded precise optogenetic interrogation, we used our second optogenetic paradigm, where inhibition was delivered for 6 s following outcomes (Fig. 7a-d). We assumed this manipulation would best reduce persistent activity and have broad effects on task engagement, even outside of light trials. We measured the total number of completed trials as a proxy for task engagement, finding that outcome suppression of PL::P-DMS pathways on 30% of trials caused a decrease in the total number of completed trials for sessions where light was used (Fig. 7c). This effect was not observed in subsequent light-off sessions (Fig. 7c), during shorter choice suppression sessions (data not shown) and could not be explained by other typical motivational regulators such as body weight (Fig. S9b). Task disengagement was also manifest as elongated initiation latencies in the PL::P-DMS outcome inhibition sessions (Fig. 7d) but was not on overall slowing of motor performance (note unchanged choice latencies in Fig. S9b). In contrast, the PL::A-DMS pathways, which exhibited weaker internal value coding, did not impact task engagement as measured by total trials or initiation latencies (Fig. 7a,b, Fig. S9a). These results suggest that temporally integrated task value signals in PL::P-DMS pathways are important for driving global task engagement.

**Figure 7.**
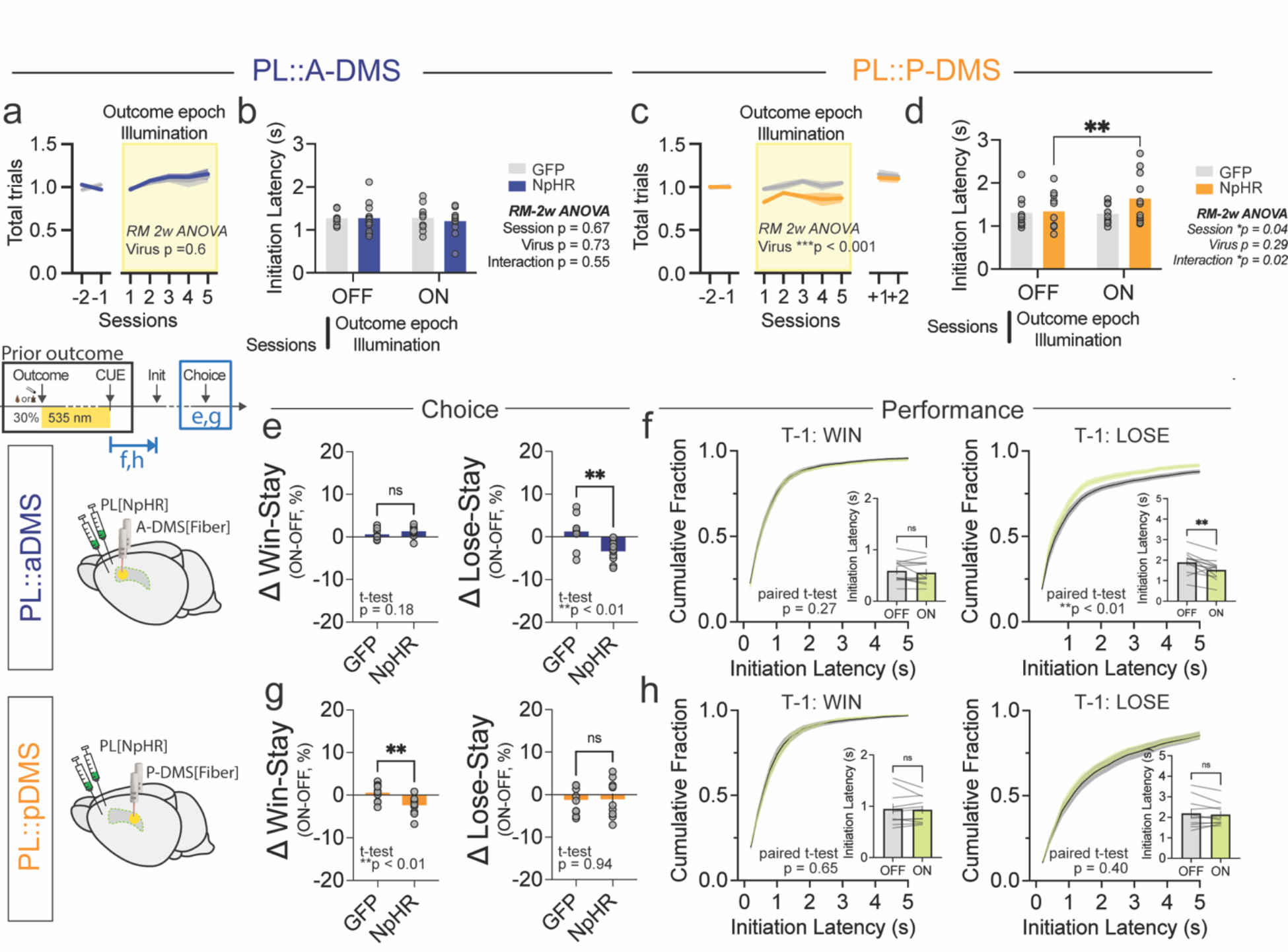
PL:: Optogenetic suppression of outcome-associated signals causes pathway- specific effects on task engagement and responses to positive and negative outcomes. a) Normalized number of total trials per session in sessions without and with random 30% outcome optogenetic inhibition of PL::A-DMS pathway (yellow bars). b) Comparison of initiation latency between sessions with (ON) or without (OFF) outcome epoch illumination of PL::A-DMS circuits from either GFP or NpHR group. c) Normalized number of total trials per session in sessions without and with random 30% outcome optogenetic inhibition of PL::P-DMS pathway (yellow bars). d) Comparison of initiation latency between sessions with (ON) or without (OFF) outcome epoch illumination of PL::P-DMS circuits from either GFP or NpHR group. e) Comparison of ΔWin- Stay(*left*) and ΔLose-Stay(*right*) between GFP and NpHR groups when light was delivered during prior trial outcomes to PL terminals in A-DMS. f) Cumulative distribution of initiation latencies and average of initiation latency (inset) following outcome light ON versus OFF trials from NpHR inhibition of PL::A-DMS. g) Comparison of ΔWin-Stay(*left*) and ΔLose-Stay(*right*) between GFP and NpHR groups when light was delivered during prior trial outcomes to PL terminals in P-DMS. h) Cumulative distribution of initiation latencies and average of initiation latency (inset) following outcome light ON versus OFF trials from NpHR inhibition of PL::P-DMS.

### PL::DMS pathways divergently control response strategies to positive and negative outcomes

To directly evaluate the divergent functions of outcome-related PL::DMS activity, we optogenetically inhibited terminals in each striatal subregion following both positive and negative outcomes (Fig. 7e-h). While we did not observe any choice or performance changes from suppression of PL::A-DMS terminals following positive outcomes (Fig.7e), we reliably observed a decrease in the win-stay probability from similar manipulations of the PL::P-DMS pathway (Fig. 7g). In contrast, we found that optogenetic suppression during negative outcomes of the PL::A- DMS, but not the PL::P-DMS, caused a robust decrease in lose-stay choice strategy (Fig. 7e,g). Furthermore, we observed similar behavioral effects for PL::A-DMS inhibition across a range of reward probability environments (Fig. S10a). Finally, we also noted that PL::A-DMS inhibition disrupted the natural slowing of trial initiations observed following negative outcomes (Fig. 7f,h)^24, 27, 28^. These results support divergent fronto-striatal control of outcome-related strategies, with PL::P-DMS activity mediating positive reinforcement and PL::A-DMS driving choice persistence in the face of negative outcomes.

### Second order retrograde tracing uncovers pathway specific afferent connectomes

Our neural coding analyses and causal manipulation studies consistently indicated a functional division of PL::DMS pathways for key neural processes that generate goal directed choice behavior. As an initial step into understanding the origins of this divergence, we examined the second-order excitatory connectomes for PL neurons defined by A-/P-DMS subregion. To do this, we injected retroAAV2-EF1a::3xFLAG-Cre into either A- or P-DMS subregions and a mixture of AAV-DJ-CAG::FLEX-TVA-mCherry and AAV-DJ-CAG::DIO-RVG into PL cortex (Fig. 8a). Subsequent PL injection of EnvA-RV-EGFP permitted single synapse tracing specifically from PL neurons that projected to either DMS subregions (2^nd^ order inputs). Consistent with these fronto- striatal circuits being embedded in the same local microcircuit, we observed multiple afferent populations with similar targeting of each PL circuit, including dorsal anterior cingulate cortex (dACC) and both associative and ventral motor thalamic nuclei (Fig. 8b,c). Surprisingly though, we also noted pathway-specific distinctions in second order afferent connections, with strong PL::P-DMS biases for secondary motor cortex (M2) and significant PL::A-DMS biases for ventral anterior cingulate cortex, retrosplenial cortex and orbitofrontal cortex. These observations suggest that the distinct coding and functional properties of PL::DMS pathways could be at least partly due to unequal strength of afferent connectivity, although other mechanisms such as divergent recurrent processing in local circuits cannot be excluded.

**Figure 8.**
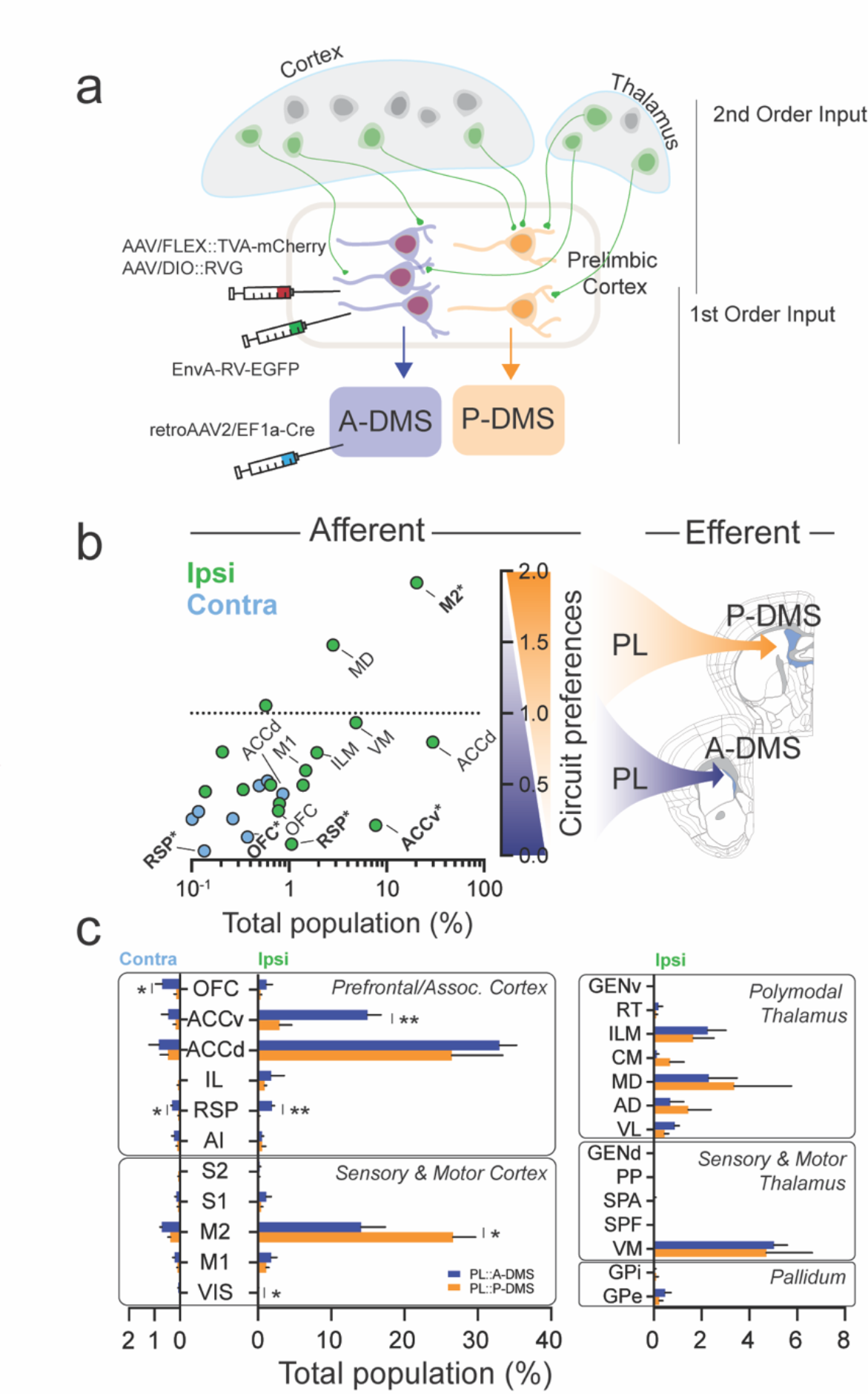
Second-order circuit tracing reveals preferential innervation of PL neurons according to A/P DMS target. a) Schematic of tracing approach to label 2^nd^ order projection to PL circuits defined by DMS target. RetroAAV2-Cre virus was injected into either A/P-DMS with Cre-sensitive TVA receptor and G-prot in PL, followed by EnvA pseudotyped-ΔG-Rabies virus one week later. b) Brain-wide innervation preferences of PL::A/P DMS pathways. Abscissa shows relative proportion (out of total labeled neurons) for each brain region and ordinate shows the ratio between pathways (PL::P-DMS/PL::A-DMS). Green and blue circles represent ipsilateral and contralateral sites relative to injectrion. Asterisks indicates statistical significance (unpaired t-test, significance *p<0.05, **p<0.01). c) Comparison of second-order innervation from major afferent brain areas.

## DISCUSSION

The dorsal striatum is a canonical set of circuits that interfaces much of the forebrain with downstream basal ganglia nuclei that select and modulate motor output^29^. Accordingly, neural processing within striatum is thought to be reflective of cortical activity^30^. Cortico-striatal projections are highly localized along the dorsal-ventral and medial-lateral axes^4^, but less so along the anterior-posterior striatal extent^3, 11^. Here we sought to understand the implications of this architecture for cortico-striatal processing, focusing on prelimbic cortical connections to dorsomedial striatum. As for most DMS-targeting afferents, we found that PL cortex formed non-overlapping circuits according to A-P target. *In vivo* imaging demonstrated that these two populations divided encoding of key behavioral variables for goal-directed choice. PL::A-DMS pathways strongly encoded choice and negative outcome, while PL::P-DMS pathways strongly encoded internal value representations and an integrated positive outcome/choice signal. Target- and temporally- specific optogenetic manipulations further confirmed the functional divergence of these fronto-striatal circuits, with PL::A-DMS pathways providing integrated responses to negative outcomes and PL::P-DMS pathways supporting task engagement and reinforcement by positive outcomes.

### Temporal and spatial distribution of goal-directed processing by PL sub-circuits

Feedback-driven goal-directed behaviors require specific response strategies to positive and negative outcomes, estimation and retention of value estimates for actions, the appropriate assignment of credit for temporally displaced choice and outcome, as well as regulation of motivation, performance, and task engagement. Here we provide evidence that prefrontal connections to the DMS supports many of these core processing functions and do so in a distributed manner across A-P striatal targets.

### Action Monitoring

Our Ca^2+^ imaging data demonstrated that PL populations projecting to the A-DMS contain neurons tuned to sensorimotor components of our operant behavioral task. While task start cue and subsequent initiation approach were represented by small subpopulations (Fig. S3a), we found that a substantial number of PL::A-DMS neurons were modulated by port choice. Averaged choice-associated kernels revealed larger contralateral than ipsilateral choice signals that occurred after choice was registered (Fig. 4e, Fig. S3c). These data are consistent with previous work showing only weak neural signals for upcoming choice in medial prefrontal cortex (mPFC), suggesting activity in this region doesn’t significantly contribute to action planning in trial and error tasks^31^. We directly probed the functional importance of choice-associated modulation via optogenetic inhibition of PL terminals within the A-DMS, finding that while bilateral optogenetic disruption of these circuits around the choice period had no effect on current trial choice selection or performance, this manipulation specifically altered choices on trials following negative outcomes (Fig. 6a). These data suggest a model where PL::A-DMS choice signals provide an efference copy of actions that is utilized to update choice values on subsequent trials. Striatal- targeting efference signals have been proposed to function together with cortical representations of environment to bind context, selected action and outcome^1, 32^. Interestingly, our choice-associated signals only seemed relevant following negative outcomes, as manipulations did not alter win-stay probabilities (Fig. 6a). These data are consistent with the biased responding of PL::A-DMS pathways towards negative outcomes (see below), suggesting common valence processing in this pathway. Recently, PL neurons that project to the nucleus accumbens core were shown to exhibit choice modulation that progressed sequentially through the population, bridging choice and outcome periods^33^. In contrast to our results, optogenetic activation of PL-NAc throughout the trial altered subsequent responses following both positive and negative outcomes.

### Outcome Monitoring

Outcome monitoring is thought to be a crucial function of prefrontal cortical circuitry, influencing how animals use subsequent sensory information^34, 35^ and select future actions^24, 27, 36^. While the PL cortex has been suggested to provide both positive and negative feedback signals to shape behavior ^37^, our experiments reveal a distribution of these functions according to DMS target, with positive outcomes encoded by both pathways and negative outcome encoding exclusively by PL::A-DMS. The PL::A-DMS pathway exhibited stronger encoding of brief (∼1s) neuronal responses to positive outcomes (Fig. S5h), while PL::P-DMS more strongly encoded positive outcomes that followed specific choices (Ch x O+ interaction; Fig. 5d). Interestingly, activity patterns for Ch x O+ coding exhibited distinct temporal patterns according to PL circuit, with a persistent (>5 s on average) activity in PL::P:DMS neurons (Fig. 5e). We hypothesized that this activity would be central to positive reinforcement behavior, either via providing an eligibility trace for plasticity or by directly influencing ensuing decision processes. To test this, we optogenetically inhibited PL::P-DMS continuously for 6 s following trial outcome, observing that stay-behavior was reduced following positive outcomes with no change in choice for manipulation following negative outcomes. These data are strongly consistent with seminal experiments showing the P-DMS to be central to outcome-driven action selection^8, 9^. Furthermore, it seems likely that this prolonged Ch x O+ activity may explain the value-based learning deficits observed upon chronic chemogenetic-mediated suppression of PL-P::DMS pathways^6^.

One surprising result of our work was the exclusive representation of negative outcomes by PL::A- DMS pathways. Averaged negative outcome kernels in this population displayed a delayed onset (∼500ms) and persistent activity lasting over 5 s, consistent with an outcome feedback signal as opposed to reward port approach (Fig. 5j). While there are numerous examples of outcome encoding in rodent PL cortex for negative valence, most cases involved aversive stimuli such as foot-shocks or air puffs^38^. A gambling task in rats, where risky maze arms had lower probability/higher reward outcomes, elicited prolonged bouts of firing in PL neurons at negative outcome that supported risky choice^39^. Choice monitoring activity was also seen at outcome in PL neurons which supported cognitive flexibility during set-shifting tasks^35^. We found that specific optogenetic inhibition of negative outcome signals in PL::A-DMS pathways reliably decreased stay behaviors following a prior loss (ie. increased choice switching), while having no choice effects following prior positive outcomes (Fig. 7e). The ability of PL::A-DMS outcome activity to support choice persistence following losses was a context-independent function, as optogenetic inhibition always decreased lose-stay behavior regardless of the probability of receiving a reward (Fig. S10). Thus, this optogenetic manipulation improved overall performance in high reward probability environments, but impaired it in lower reward probability scenarios, where lose-stay behavior is adaptive (not shown). This data argues against a role for mPFC circuits in flexibly supporting behavioral strategies following negative outcome. Furthermore, these functional effects strongly contrast with negative outcome-tuned neurons in the ACC, which have been shown to implement choice switching in many species^40, 41^. While response persistence in the face of negative outcomes is essential in sparse reward environments, left unchecked this tendency could clearly impair value-based function. This raises the question of whether mouse models of neuropsychiatric disease characterized by perseverative choice abnormalities exhibit dysregulation of PL::A-DMS pathways.

### Internal Representations of Value

Internal representation of choice value and local reward availability are key determinants of behavior in dynamic foraging tasks^24, 27^. Our results suggest that PL::P-DMS pathways more strongly encode these behavioral parameters as compared with PL::A-DMS pathways. We found that relative value signals tracked strongly with the baseline, but not phasic components of cellular calcium signals (Fig. 4g,l, Fig. S4e,f). Our ΔQ-encoding PL population is consistent with a previously identified PL-DMS population that stably represented relative value via persistent baseline spiking activity^24^. While we also identified neural signals encoding total choice value (*Σ*Q) as in Bari et al., our inability to control trial initiation precluded investigation into the relative persistence of these distinct value signals^24^. Engagement in self-initiated foraging tasks is strongly modulated by local reward environment, a variable we captured with a local average of the reward rate. Again, we found that PL::P-DMS pathways more strongly encoded this feature as compared to PL::A-DMS pathways. The persistent nature of value coding in these pathways made phasic optogenetic manipulation difficult. To circumvent this, we looked at all trials in sessions where inhibition was delivered in 30% of trials for 6 s after outcome. We reasoned that prolonged inhibition should sufficiently alter persistent neural signals to impact immediately subsequent trials as well as the overall behavior of the animal in the session. Indeed, we found that post-outcome inhibition was able to both reduce the total number of initiated trials and reduce the win-stay probability in non-light trials, suggesting the involvement of reward-rate and ΔQ-encoding PL::P- DMS populations, respectively. It is interesting to hypothesize that the reduction in task engagement caused by disruption of this pathway may share a common cause with the reduced responding seen in earlier P-DMS lesion studies^8^.

### What underlies the functional diversification of PL cortex?

Our work adds to recent studies demonstrating a range of behavioral functions for PL cortical microcircuits defined by target area^20–22, 33, 35^. Nevertheless, the mechanisms underlying this functional diversification remain unclear, with potential candidates including circuit-specific differences in molecular composition, long-range afferent projections, or local synaptic networks. While evidence exists for target-specific transcriptional differences in PL cortex^20^, other analyses have shown diverse PL functions emerging from molecularly homogenous populations^22^. Circuit- specific transcriptional profiling could reveal whether molecular diversity can account for divergent PL-DMS pathway activity. Differences in afferent connectivity may result from circuit-specific differences in local inhibitory control^42^ or long-range excitatory projections. We used 2-stage retrograde tracing to map afferent populations that synapsed on PL neurons defined by A/P-DMS target (Fig. 8), finding that ACCv, RSP cortex and OFC were strongly biased towards PL::A-DMS populations while M2 connectivity favored PL::P-DMS. Upstream manipulations will be necessary to test whether prolonged choice encoding in M2^36^ supports persistent Ch x O+ signaling in PL::P- DMS neurons, while enhanced ACCv, RSP and OFC connectivity to PL::A-DMS supports negative outcome associated activity. Similar tracing approaches have highlighted the importance of ACCv connectivity to deep PL layers projecting to NAc for outcome monitoring during cognitive flexibility tasks^35^.

### Functional implications of this circuit architecture

Our initial tracing data showed a surprising number of cortical and thalamic regions have distinct, yet intermingled populations projecting to A/P-DMS (Fig. S1). Future work should explore the computational advantages afforded by this arrangement. It is presently unclear whether anterior and posterior striatal subregions might work coordinately or antagonistically to control behavior, which would be an important starting point for our understanding. Either way, this organization could permit appropriate and flexible coordination of A/P-DMS targeting populations via local-circuit interactions in cortex or thalamus. Alternatively, these parallel processing paths may be integrated via downstream basal ganglia components.

## Acknowledgement

This work was supported by R01MH115030 to M.V.F and Whitehall Foundation grant to M.V.F. We thank Dr. Long Ding, Dr. Elizabeth N. Holly, Dr. Mariexcel F. Davatolhagh for useful feedback on the manuscript. We also thank Alexandria J. Cowell, Alessandro Jean-Louis, Sara Seyedroudbari, Michaela Glass, John Talley, Aaron Uy for providing assist.

## Author Contributions Statement

Conceptualization, K. C. and M.V.F.; methodology, K.C., E.P. and M.V.F.; formal analysis, K.C., E.P., L.V., N.T.H. and E.D.; investigation, K.C., writing – original draft, K.C. and M.V.F.; writing – review and editing, K.C., E.P., and M.V.F.; visualization, K.C.; supervision, K.C., E.P., C.R.G. and M.V.F.; Funding Acquisition, M.V.F.

## METHODS

### Animal

Animal experiment procedures were approved by the *University of Pennsylvania Institutional Animal Care and Use Committee*, and all experiments were conducted in accordance with the *National Institutes of Health Guidelines for the Use of Animals*. All animals were supplied from Charles River Laboratory (Wilmington, MA, strain code 027, C57BL/6NCrl). Unless otherwise noted, animals were grouped with littermates on a 12:12 light-dark cycle and provided *ad libitum* food and water. All experiments were conducted on naive male mice.

### Stereotaxic surgery

Intracranial surgery was conducted on a stereotaxic surgery frame (Kopf Instrument, Model 1900) under isoflurane anesthesia (1.5-2% + oxygen 1 L/min). Animal body temperature was maintained at 30°C during surgery using a feedback thermocontroller (Harvard apparatus, #50722F). Skin was cleaned with Nair hair remover followed by application of betadine to disinfect the area. Prior to surgery, 2mg/kg bupivacaine was administered subcutaneously, and the mouse was given a single dose of meloxicam (5mg/kg). Skin was carefully opened along the anterior-posterior midline, bregma was set to zero based on skull balance. A craniotomy was performed with a drill above the target site. Virus or Tracer was loaded into mineral oil (Sigma-Aldrich, M3516)-filled glass pipette (WPI, TW100F-3) and delivered at rate 30 nl/min using a micro-infusion pump (Harvard Apparatus, #70-3007). Pipette was carefully withdrawn from the brain, and the skin was sutured. Animals were monitored up to 1 hour following regaining of consciousness, then transferred to the home cage and monitored after 24h, 48h and 72h. Injection coordinates, A-DMS: AP + 1.2 mm, ML +1.35 mm, DV -2.7 mm; P-DMS: AP -0.3 mm, ML + 1.95 mm, DV -2.2 mm; PL: AP + 2.0 mm, ML +0.35 mm, DV -1.7 mm

### Anatomical Tracing

For mapping PL synaptic terminals in DMS (anterograde tracing), a 1:1 mixture of AAV5- CaMKii::Cre + AAVdj-EF1a::Flex-Synaptophysin-mRuby viruses were injected into PL. For mapping retrogradely labeled neurons, a mixture of AAV1-Syn::Cre + AAVdj-EF1a::DIO-RVG + AAVdj-EF1a::Flex-TVA-mCherry was injected into A-DMS and Alexa647-conjugated Cholera toxin subunit-B was injected into P-DMS. Seven days later, EnvA G-Deleted Rabies-eGFP was injected into A-DMS. For mapping 2nd-tier projections to PL::DMS pathway, retroAAV2- hSyn::3xFlag-Cre was injected into the A- or P-DMS followed by a mixture of AAVdj-EF1a::DIO-RVG + AAVdj-EF1a::Flex-TVA-mCherry into PL. Seven days later, EnvA G-Deleted Rabies-eGFP was injected into PL.

After viral injection, 7 days (for Rabies virus) or 3 weeks (for AAV) were allowed for viral expression, animals were deeply anaesthetized with i.p injection of 100 uL pentobarbital sodium (Nembutal, 50 mg/mL) and transcardially perfused with PBS followed by formalin (10%). Brains were removed and post-fixed in formalin (Thermo Fisher Scientific, SF1004) overnight, then transferred to PBS. Brains were sectioned coronally at 50µm then brain slices were mounted on slide glasses and covered with fluoromount solution (SouthernBiotech, #0100-01) for imaging.

Stitched large-field images were obtained with a 4x objective (Olympus, 4x, 0.16NA) on an epi- fluorescent microscope (Olympus, BX63). Fluorescence-positive neurons were counted using automated object detection (NeuroInfo Suite, v2021.). Three-dimensional brain images were then reconstructed using NeuroInfo software (MBF bioscience), which registered individual slices to the Allen Institute reference brain atlas (Allen mouse common coordinate framework; CCFv3) ^43^.

### Immunohistochemistry

At room temperature, free floating brain slices were permeabilized in 0.6% Triton x-100 and blocked with 6% normal goat serum (Jackson ImmunoResearch, 005-000-121) in PBS for 1 h. Samples were incubated in primary antibody solution (Rat anti-CTIP2, 1:500, Abcam, ab18465) overnight in 0.2% Triton x-100 and 2% NGS in PBS. Slices were washed then incubated in secondary antibody solution (Goat anti-rat IgG-alexa555 conjugated, 1:500, Invitrogen, A48263) for 1h in 0.2% Triton x-100 and 2% NGS in PBS, then mounted and imaged.

### Behavioral equipment

Behavior training was conducted utilizing a custom built 3-port operant chamber (dimensions 7.5 L x 5.5 W x 5.13 H inches, Sanworks LLC, NY). Each port is controlled by a TTL signal from the state machine consisting of white LED light, infrared beam break detector and liquid outlet. The center port was designated as a reward delivery outlet using a pinch valve (225P011-21, NResearch, NJ). All behavior chambers were enclosed in sound-attenuating boxes (PSIB27, Pyle, NY). Behavior protocols were controlled by Bpod software (https://github.com/sanworks/Bpod) in MATLAB (MathWorks). All port entries and events were recorded by the Bpod State Machine during behavioral sessions.

### Behavioral Training

To increase operant responding, total calorie consumption was reduced over 1 week to reach 85∼90% body weight, a level maintained throughout operant training. Animals were habituated to behavior chambers for at least 2-days prior to training. Each day, animals were given 45 min of exposure to the behavioral box with chocolate milk (Boost Original ready to drink, rich chocolate nutritional shake, Nestle) delivery from the center port spaced 20 s. apart. Following the habituation period, animals performed light-guided sessions as follows: 1) center port light indicated the beginning of a trial; 2) trial initiation via a center poke led to illumination of a randomly selected side port; 3) appropriate selection of the lit port within 3 s led to illumination of the center port and delivery of 12ul of Boost at this location; 4) selection of the unlit alternative led to illumination of the center port without concomitant dispensing of reward. Each trial was separated by a 5 s. ITI in which all chamber lights were extinguished. Sessions lasted 1 hour with no trial limits. Animals were considered to reach criteria with >200 completed trials per session.

### Two-alternative forced choice task

After reaching criteria performance levels in light-guided training, mice progressed to a 2- alternative forced choice task structured as follows: 1) center port light indicated the beginning of a trial; 2) a 500 ms. holding period (sequentially increased from 0ms, 100ms, 300ms) in the center port triggered the illumination of both side ports; 3) animals had a 3 s. window to register either left or right port choice. When animals failed to make a choice in this period this resulted in an omission, which was followed by a 3 s timeout and required the animal to reinitiate the trial. 4) successful registration of a choice was followed by 0.5 s delay period ending in the outcome period (*P*_outcome_ = 85%). Correct choice resulted in delivering 12μL Boost from the center port with 85% chance while incorrect choice resulted a in brief punishment tone (white noise) with 3 s timeouts, also with 85% chance. In the remaining 15% of trials, animals didn’t receive any outcome (reward or punishment). Each trial was separated by a 3 s. ITI in which all chamber lights were extinguished. To prevent outcome-insensitive behavior, past-reward history was monitored in a 10-trial moving window and rewarded side was switched when 8 of the last 10 choices were to the currently rewarded port. Sessions lasted for either 45 min. (1-p imaging) or 1 hr (optogenetic manipulations). We utilized a relative reward-stay value >2 to decide when to move mice on to recording sessions. Relative reward stay was defined as:

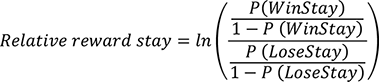

### 1-p Imaging

To record calcium signals from PL neurons targeting specific striatal subregions, retroAAV2/EF1a-3xFlag-Cre^44^ was unilaterally injected to A- or P-DMS, together with AAV1/hSyn-Flex::GCaMP7f-WPRE^45^ injection into PL. Prior to relay GRIN lens (1mm x 4mm, Inscopix, 1050- 002176) implantation in PL, upper prefrontal tissue was gently aspirated using a glass pipette until reaching 0.5 mm above target site. Following tissue aspiration, the GRIN lens was slowly lowered (100μm/min) until 0.3 mm above from the target site. Dental cement (Geristore™) was used to create a foundation around the GRIN lens, and the remaining exposed GRIN lens was covered with silicone paste to prevent scratches. After surgery, animals were transferred to a single housed cage, where their status was monitored until movement recovery. The anti-inflammatory Meloxicam (5 mg/kg) was applied subcutaneously daily for >1 week, and animals were carefully monitored. Four to six weeks following GRIN lens implantation, the miniscope baseplate was installed under 1-p imaging (UCLA miniscope v3.0)^46^ to locate fields of view (FOV) with robust GCaMP7f expression. Once a FOV was selected, the baseplate was fixed with dental cement to make a crown. Baseplates were covered with a cap, and animals were subsequently returned to the home cage.

### Signal processing

To extract calcium traces from imaging videos, we utilized MIN1PIPE (v2 alpha, https://github.com/JinghaoLu/MIN1PIPE/tree/v2-alpha) for motion correction (Hierarchical non-rigid movement correction), segmentation (GMM, LSTM classifier), and signal deconvolution (CNMF identifier)^47^. Each ROI selected by MIN1PIPE was individually reviewed to ensure somatic morphology and remove repeated selection of the same neuron’s proximal dendrites.

### Neural Encoding Model

To analyze task-relevant neural activity we designed a neural encoding model based upon a linear combination of event-based and continuous predictors. We included episodic external variables, including trial start cue, self-initiation, choice and outcome, which were represented by spline-based, temporally expanded kernels (Fig. 2d). In addition, we sought to identify neural signals encoding relevant internal value information likely guiding choice^48^. To do this we fit our choice data with a Q-learning reinforcement model and used Q values and reward prediction errors as internal behavioral variables. Specifically, we included trial-by-trial ΣQ (Q_left_ + Q_right_) and ΔQ (Q_left_- Q_right_) values (as continuous predictors that changed their value at outcome, consistent with the RL model) and reward prediction errors (kernels tethered to the outcome). Finally, we included a local reward rate averaged over the prior 5 trials as a continuous behavioral variable. We fit these regression parameters using a generalized linear model with near-lasso (elastic net, alpha=0.95) regularization to achieve sparse regression weights, using the *glmnet* library^49^ (wrapped for MATLAB usage with custom software available at doi:10.5281/zenodo.3568314). Details on the representation of each predictor in the design matrix of the model are given in Table 1. For each fitted trace, the shrinkage hyperparameter λ controlling the strength of the elastic net regularization was selected by 50-fold cross-validation. Following established practice (Hastie, 2008), given the maximum value of the (cross-validated) fraction of variance explained (FVE) over possible values of λ and its standard deviation across folds, we selected the largest value of λ such that its associated FVE was larger than the maximum minus one standard deviation, thus selecting the “simplest” model in the neighborhood of the best-fitting one. The cross-validation folds were stratified by experimental trial, so that each trial was represented roughly equally in each fold, and the data was grouped in 200ms-long temporal chunks (typically corresponding to about 4 imaging frames) for the purpose of cross-validation, to reduce the number of temporally adjacent data samples in different cross-validation folds^50^.

We fitted an independent model to each recorded neuron. Using the model, the fraction of variance explained (FVE) was calculated by comparing the full model and actual calcium signal trace as an indicator of the accuracy of model prediction. To exclude non-task relevant neurons, we limited further analyses to those with at least 5% FVE. To assess contribution from a certain predictor, we calculated a tuning index defined as FVE(Full) – FVE (reduced), where FVE (reduced) is the FVE of the reduced model obtained by removing the predictor of interest from the full model. Tuning to a group of predictors was quantified in the same way. This tuning index gives a lower bound on the amount of variability in the data that can be explained by the predictor (or group of predictors) of interest. Whenever a dichotomous “tuned”/“not Tuned” characterization was needed, such as in the donut plots in Fig 3, we classified as “tuned” neurons that met a 5% tuning threshold for grouped predictors (Fig 3d).

Each neuron’s fitted model provided tuning estimates for all predictors, as well as neuron-level estimates of the model kernels such as those in Fig 4a, 5a, f. The pathway-level kernels in Fig 4e, 5e,j were defined as the root-mean-square of the kernels of task-relevant neurons in each pathway, and their confidence intervals were determined by bootstrapping over the set of neurons (10,000 bootstrap samples). The pathway kernel for a certain predictor can be given an intuitive geometrical interpretation as follows: if we consider the pseudo-population vector describing the activity of all recorded (and task-relevant) neurons within a pathway, the pathway kernel for a predictor at lag *t* is an estimate of the distance at lag *t* of the population vector from its time-averaged value, after accounting for the effect of the other predictors. By construction, then, the pathway kernels can never be negative, as they capture the overall magnitude of the effect of the predictor on the neural population, rather than a specific modulation direction. The statistical significance of the difference of pathway kernels in Fig 4c, 5dh was assessed with a bootstrap test ^51^, performed with the Bias-Corrected and accelerated(BCa) technique and bootstrapping over the set of neurons belonging to the two pathways (10,000 bootstrap samples).

### Reinforcement Learning Model

We adapted a Q-learning Reinforcement Learning Model with two basic parameters that fit the behavioral data produced by the serial reversal task. Mouse choice and outcome history were the primary inputs of the model. The values of the choice alternatives were initiated at 0 and updated as follows:

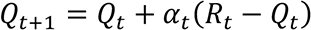

Qt is the value of the action taken on trial t of each choice and R is the actual reward received in trial t. Learning rate was controlled by the parameter α. Softmax rule was employed to infer trial-by-trial Q-values for each choice, which relates choice probability to differences in choice value: 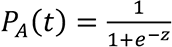, where

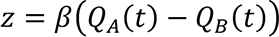

*β* is the inverse temperature parameter and controls the degree to which choices are biased by perceived value. High values for *β* indicate mice more readily exploit differences in action values between the alternatives, while lower values suggest that mice exhibit more exploratory behavior. To fit this model to our choice data, we used the *fmincon* function in MATLAB to minimize the negative log likelihood of models using our parameters (*α, β*).

To quantify the goodness of fit of the RL model in an easily interpretable way, we computed the fraction of deviance of the choice data explained by the model (FDE) ^52^. The deviance of the RL model is

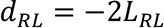

where *L* is the log-likelihood of the fitted model:

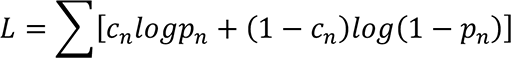

where *c_n_* is the choice on trial *n* (*c_n_*=0 for left choice, *c_n_*=1 for right choice), and *p_n_* is the probability that the model assigns to choosing right on trial *n*. The FDE is defined as

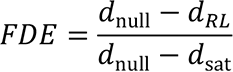

where *d*_null_ is the deviance for a null model that assigns the same probability *p_n_*=*p* on each trial (equal to the average probability of choosing right over the session), and *d*_sat_ is the deviance of the “saturated” model, which assigns *p*_n_=1 on trials where the mouse chose right, and vice versa p_n_=0 on trials where the mouse chose left. By inspection of the definition of *L* we find that *d*_sat_ is always zero, and therefore

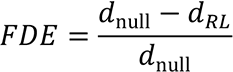

We note that FDE≤1, with FDE=0 for a model that doesn’t do better than chance (i.e., making always the same prediction for each trial) and FDE=1 for a model that explains all variability in the data (this is in general unachievable on unseen data, unless the process generating the data is deterministic). Applying this metric showed that the reinforcement learning model successfully captured the main behavioral patterns exhibited by the animals, explaining overall 23.4% of the deviance (as measured on the training data; Fig. S2c).

As a further check on the performance of the RL model, we compared its goodness of fit and predictive power with that of a logistic regression model that predicted the upcoming choice based on the choices and outcomes of the previous 5 trials. The two models had largely comparable performance (Fig S2c). The logistic regression model tended to have slightly higher FDE on the training data (Fig S2c), but this was likely an overfitting effect due to the larger number of parameters available to that model (11 parameters for the logistic regression vs 2 parameters for the RL model), as evidenced by the similar values of the Akaike Information Criterion between the two models (FigS2c).

### Optogenetics

To evaluate the behavioral role of each pathway (PL::A-DMS or PL::P-DMS), we used a Halorhodopsin-induced terminal suppression strategy (Felix-Ortiz et al., 2013). AAV5/CaMKii- NpHR3.0-eYFP (UNC vector core) or AAV5/Syn-GFP (Penn vector core) was injected bilaterally to the PL followed by bilateral implantation of custom-made optic cannulas (Thorlabs, FT200EMP, SFLC230) for pathway-specific light delivery into either A-DMS and P-DMS. To ensure full expression of NpHR in the axonal terminal, a recovery period of at least 5 weeks was allowed after viral injection. Animals were acclimated to the fiber optic tethers for at least 5 days before any behavioral sessions. Once animals performed >200 trials/day with a relative reward stay >2 we proceeded to the optogenetic manipulation phase, the training proceeded to the light delivery stage. In the behavior task, two light delivery protocols (∼530nm light from either PrizmatixFC- LED-535-TR or Shanghai DPSS, SDL-532-100T) were used to assess temporally distinct contributions from each pathway. To prevent light-induced non-specific effect, light intensity was adjusted to 0.8∼2 mW at the fiber end^53^. Choice epoch manipulations were continuous illumination from initiation poke to the end of the reward delivery delay following choice. Outcome epoch manipulations were continuous illumination from outcome delivery until next trial center light on. Either *Δ*Win-Stay or *Δ*Lose-Stay was calculated as:

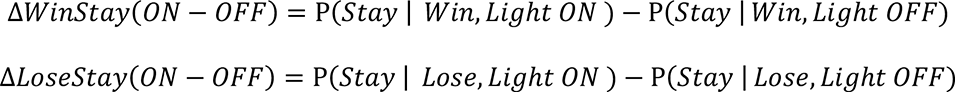

where Win or Lose indicates reward history on prior trial. Light On/Off refers to presence or absence of light illumination on previous choice epoch (Fig. 6, Fig. S8), current choice epoch (Fig. S8) and previous outcome epoch (Fig. 7, Fig. S9). Behavioral data were collected for multiple days to obtain enough trials (3351 ± 143 trials, mean ± s.e.m. across animals). Unless otherwise noted, probability of reward was 85%. For Fig. S10, some PL::A-DMS outcome optogenetic sessions were performed with reward probabilities of 1.0 or 0.4, applied to both ports.

### Quantification and statistical analysis

All data were analyzed with prism8.0 and custom MATLAB code, available upon request. Repeated measures ANOVA t-test (paired and unpaired) were performed using Prism 8.0 built-in-function. K-S tests were performed as indicated in results using *kstest2* functions in MATLAB. Kernel density estimates were performed as indicated in results using *ksdensity* functions in MATLAB. Significant effects and p-values are indicated in the figures and legends.

---

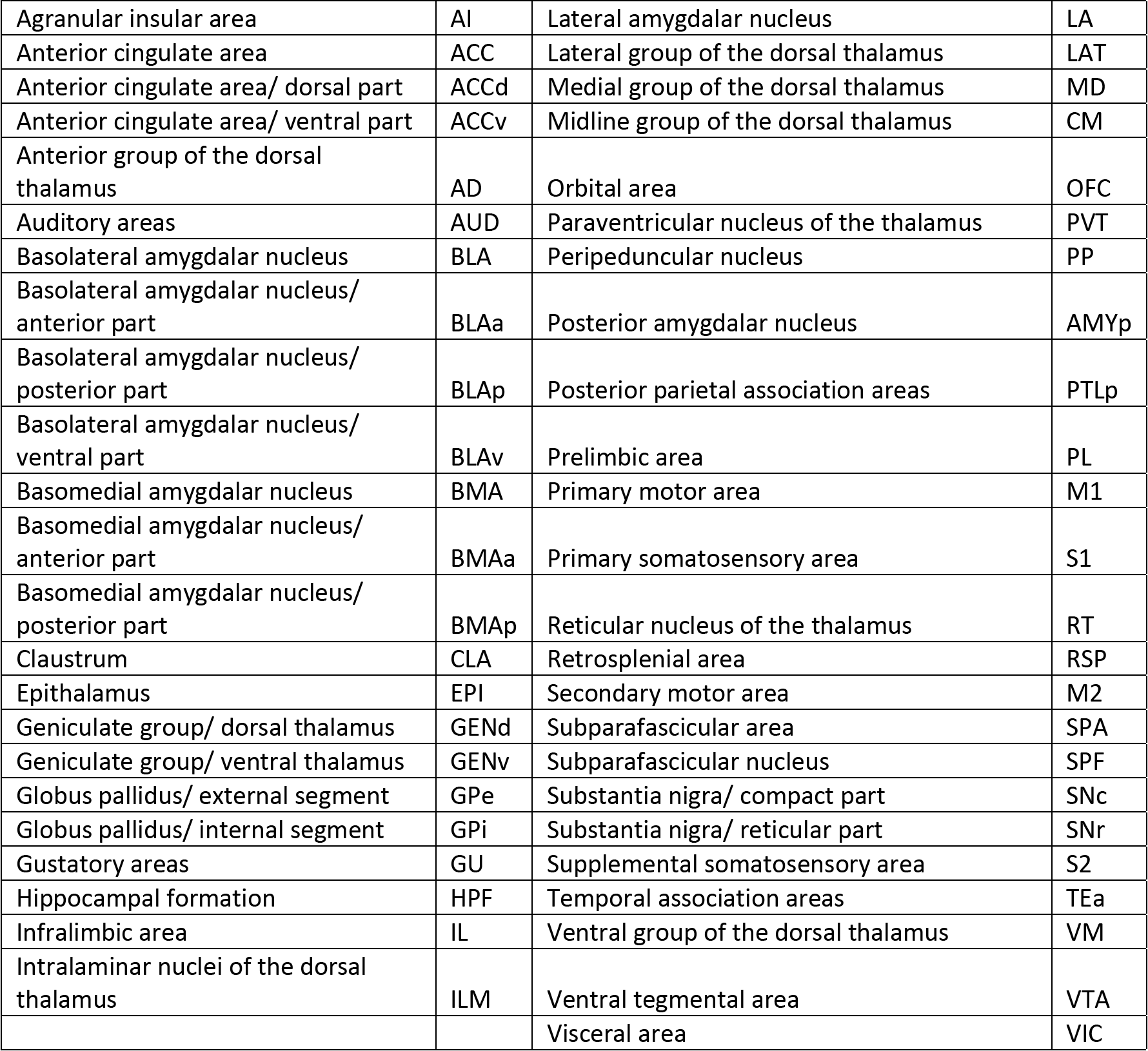
Abbreviation.

Abbreviation
|  |  |  |  |
| --- | --- | --- | --- |
| Agranular insular area | AI | Lateral amygdalar nucleus | LA |
| Anterior cingulate area | ACC | Lateral group of the dorsal thalamus | LAT |
| Anterior cingulate area/ dorsal part | ACCd | Medial group of the dorsal thalamus | MD |
| Anterior cingulate area/ ventral part | ACCv | Midline group of the dorsal thalamus | CM |
| Anterior group of the dorsal thalamus | AD | Orbital area | OFC |
| Auditory areas | AUD | Paraventricular nucleus of the thalamus | PVT |
| Basolateral amygdalar nucleus | BLA | Peripeduncular nucleus | PP |
| Basolateral amygdalar nucleus/ anterior part | BLAa | Posterior amygdalar nucleus | AMYp |
| Basolateral amygdalar nucleus/ posterior part | BLAp | Posterior parietal association areas | PTLp |
| Basolateral amygdalar nucleus/ ventral part | BLAv | Prelimbic area | PL |
| Basomedial amygdalar nucleus | BMA | Primary motor area | M1 |
| Basomedial amygdalar nucleus/ anterior part | BMAa | Primary somatosensory area | S1 |
| Basomedial amygdalar nucleus/ posterior part | BMAp | Reticular nucleus of the thalamus | RT |
| Clastrum | CLA | Retrosplenial area | RSP |
| Epithalamus | EPI | Secondary motor area | M2 |
| Geniculate group/ dorsal thalamus | GENd | Subparafascicular area | SPA |
| Geniculate group/ ventral thalamus | GENv | Subparafascicular nucleus | SPF |
| Globus pallidus/ external segment | GPe | Substantia nigra/ compact part | SNc |
| Globus pallidus/ internal segment | GPi | Substantia nigra/ reticular part | SNr |
| Gustatory areas | GU | Supplemental somatosensory area | S2 |
| Hippocampal formation | HPF | Temporal association areas | TEa |
| Infralimbic area | IL | Ventral group of the dorsal thalamus | VM |
| Intralaminar nuclei of the dorsal thalamus | ILM | Ventral tegmental area | VTA |
|  |  | Visceral area | VIC |

