## Supplemental Figures for "Distributed processing for action control by prelimbic circuits targeting anterior-posterior dorsal striatal subregions"

### supplemental Figure1

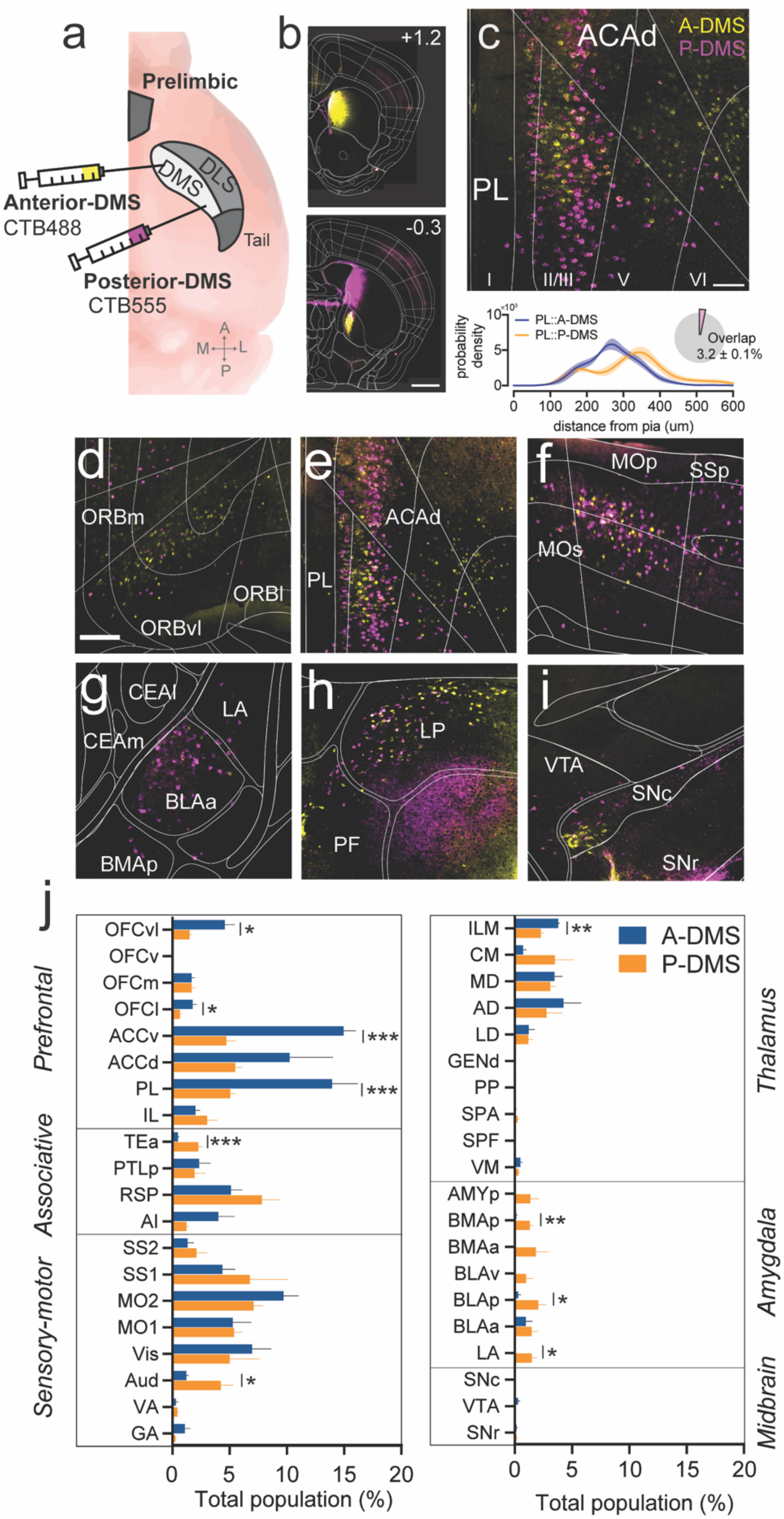

**Fig. S1. Assessment of brain-wide afferent inputs to anterior/posterior DMS compartments.** a) Schematic showing dual retrograde tracing strategy using both Alexa488 conjugated-CTB in A-DMS (yellow) and Alexa647-conjugated CTB in P-DMS (magenta). b) Example coronal section showing injection sites (*top*: A-DMS, *bottom*: P-DMS). Scale bar, 500 μm. Numbers in upper right corners indicate A/P coordinate from bregma. c) Representative image from prelimbic coronal section (*top*) and quantification of neuronal distribution from the pia (*bottom*) and overlapping population (inset). scale bar, 100 μm (n= 4). d-i) Example coronal sections of major sources of afferent inputs to A/P DMS. j) Quantification of relative proportion of labeled neurons targeting A/P DMS (unpaired t-test, significance $*p<0.05$, $**p<0.01$, $***p<0.0001$ ).

### Supplement Figure2.

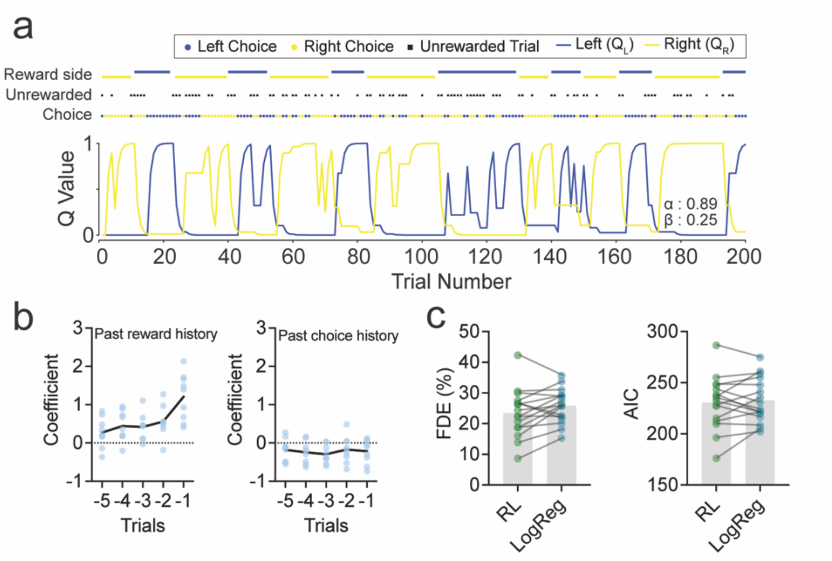


**Fig. S2. Modeling of choice behavior in value-based task.** a) Trial-by-trial choices, outcomes and predicted Q-values *(blue*, $Q_L$; *yellow*, $Q_R$) derived from Q-learning reinforcement model. b) Logistic regression model of behavior, using reward/choice history of past 5 trials as predictors. c) Fraction of deviance explained (FDE, *left*) and Akaike Information Criterion (AIC, *right*) for the logistic regression model (LogReg) and reinforcement learning model (RL) of mouse choice data. See Methods for details.

Supplemental_Fig3

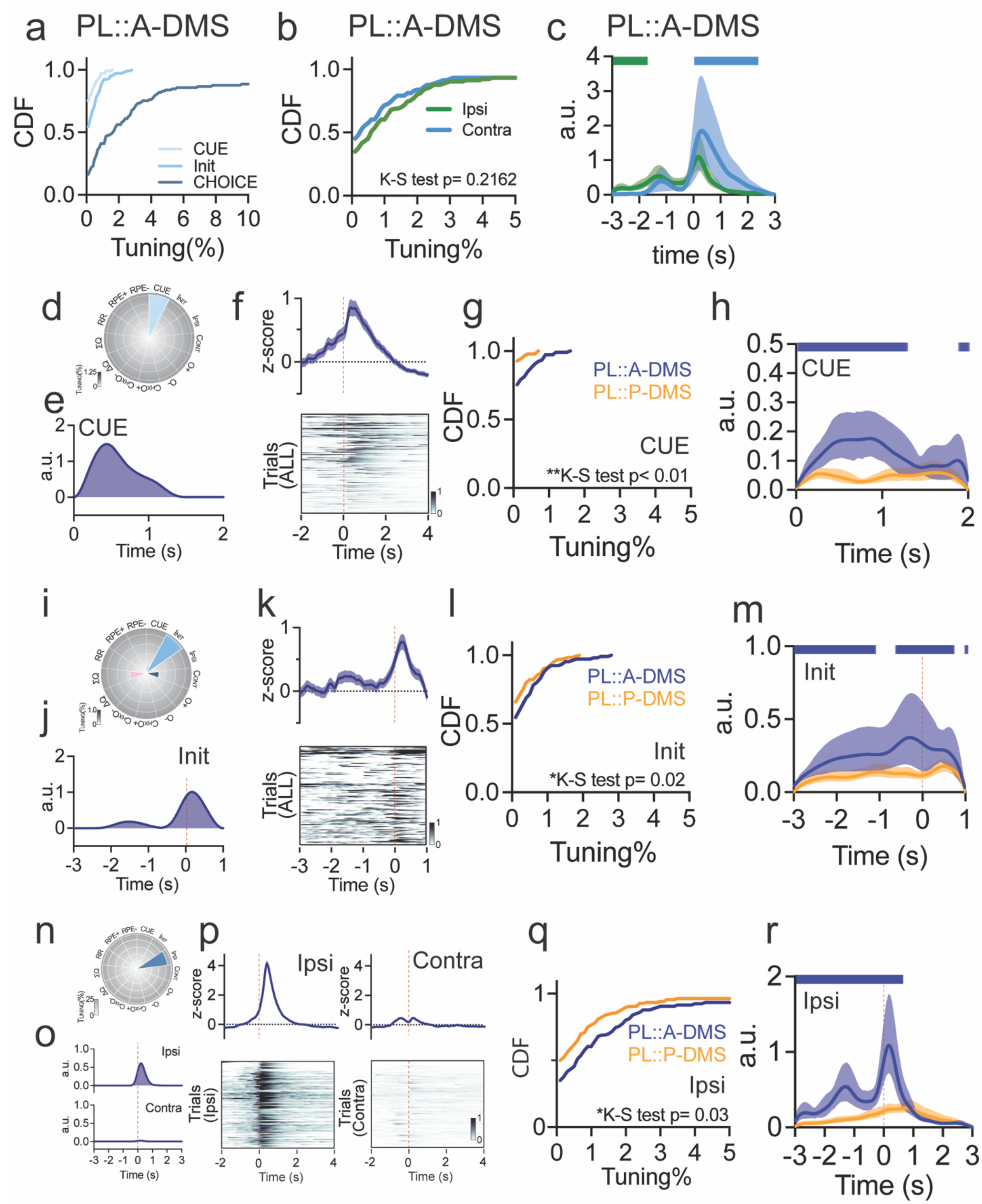

**Fig. S3. PL::A-DMS is strongly tuned to choice and weakly tuned to cue and task initiation.** a) Cumulative distributions of individual external predictors from PL::A-DMS. b) Cumulative distributions of ipsi/contra choice predictors from PL::A-DMS. c) Averaged ipsi/contra choice kernels for PL::A-DMS circuit. d) Tuning plot of CUE tuned neuron. e) Inferred CUE kernel. f) z-scored PETH (*top*) and trial-by-trial normalized neuronal activity (*bottom)* aligned to CUE. g) Cumulative distributions for CUE tuning from task-tuned neurons of both PL::DMS pathways. h) Pathway comparison of temporal dynamics for CUE modulation. i) Tuning plot of initiation tuned neuron. j) Inferred initiation kernel. k) z-scored PETH (*top*) and trial-by-trial normalized neuronal activity (*bottom)* aligned to Init. l) Cumulative distributions for Init tuning from task-tuned neurons of both PL::DMS pathways. m) Pathway comparison of temporal dynamics for initiation modulation. n) Tuning plot for ipsi. choice tuned neuron. o) Inferred kernels corresponding to Ipsi (*top*) and Contra choice (*bottom*). p) z-scored PETH (*top*) and trial-by-trial normalized neuronal activity (*bottom*) corresponding to Ipsi (*left)* and Contra choice (*right*). q) Cumulative distributions for Ipsi tuning from task-tuned neurons of both pathways. r) Pathway level of temporal dynamics for modulation aligned to Ipsi. choice.

### Supplemental Figure 4

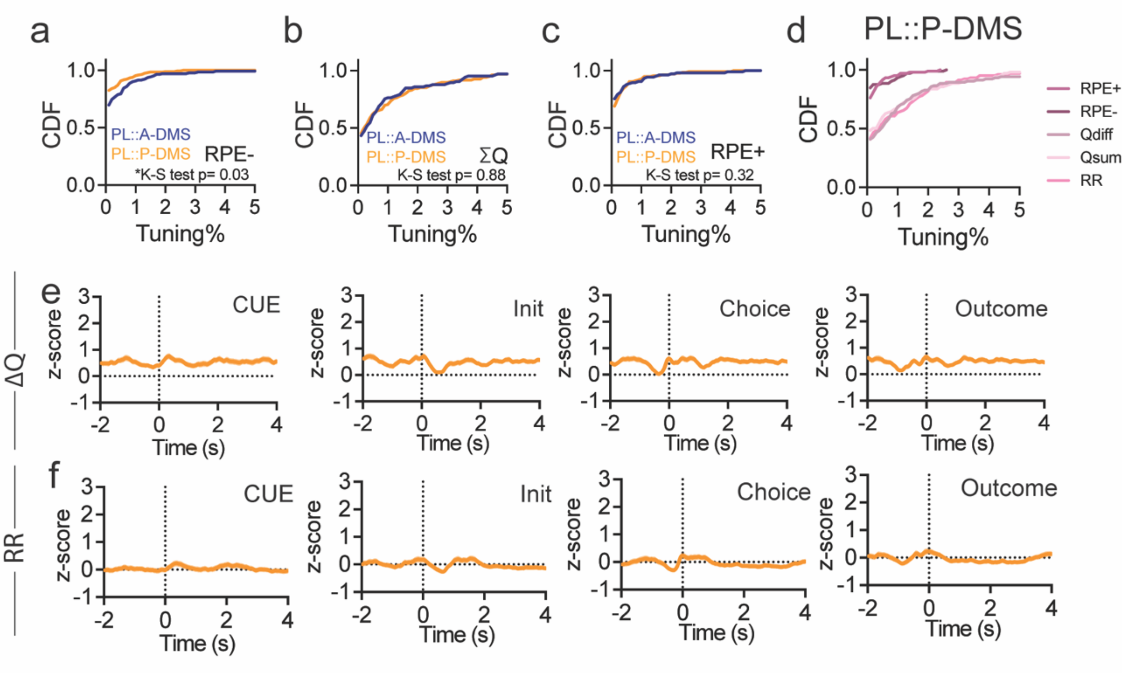


**Fig. S4. Detailed analysis of internal value neural coding.** a-c) Pathway comparison using cumulative distributions of (a) RPE-, (b) ∑Q, (c) RPE+ tuning from task-tuned neurons. d) Cumulative distributions for individual internal predictors from PL::P-DMS. e) z-scored PETH for ∆Q encoding neuron corresponding to Fig. 4f, aligned to behavioral events in right corner. f) z-scored PETH for RR encoding neuron corresponding to Fig. 4k, aligned to behavioral events in right corner.

### Supplemental Figure 5

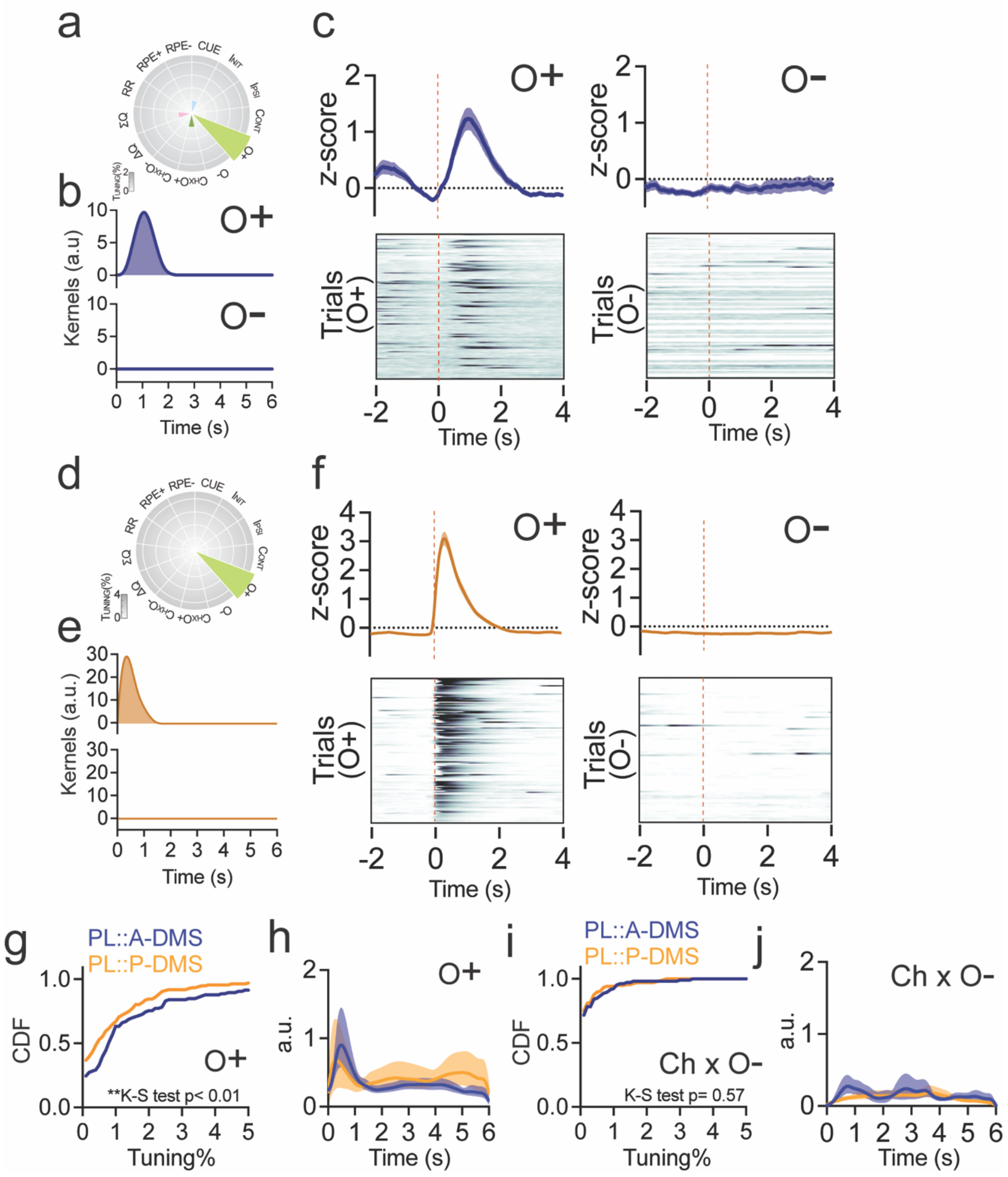


**Fig. S5. Detailed analysis of outcome encoding.** a) Tuning plot of O+ neuron from PL::A-DMS. b) Inferred kernels corresponding to O+ (*top*) and O- (*bottom*) from PL::A-DMS. c) z-scored PETH (*top*) and trial-by-trial normalized neuronal activity (*bottom*) corresponding to O+ (*left)* and O-(*right*) from PL::A-DMS. d) Tuning plot of O+ neuron from PL::P-DMS. e) Inferred kernels corresponding to O+ (*top*) and O-(*bottom*) from PL::P-DMS. f) z-scored PETH (*top*) and trial-by-trial normalized neuronal activity (*bottom*) corresponding to O+ (*left)* and O-(*right*) from PL::P-DMS. g) Pathway comparison using cumulative distributions for O+ tuning from task-tuned neurons. h) Pathway comparison of temporal dynamics for O+ modulation. i) Pathway comparison using cumulative distributions for Ch x O- tuning from task-tuned neurons. j) Pathway comparison of temporal dynamics for modulation corresponding to Ch x O-.

### Supplemental Figure 6

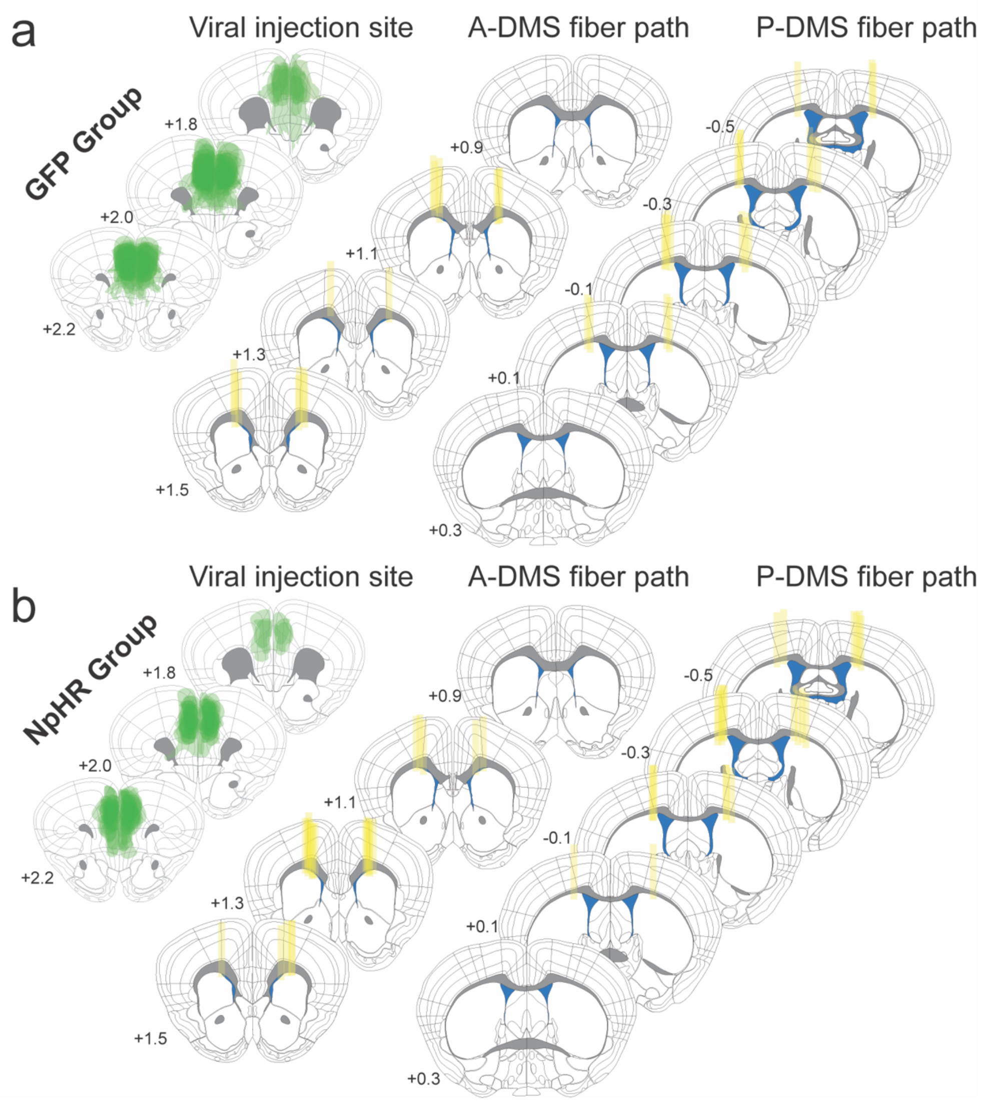


**Fig. S6. Injection sites for optogenetic experiments.** a) Brain atlas showing GFP virus injection site (left) and bilateral fiber implant site from A-DMS (center) and P-DMS groups (right). b) Brain atlas showing NpHR Virus injection site (left) and bilateral fiber implant site from A-DMS (center) and P-DMS groups (right).

### Supplemental Figure 7

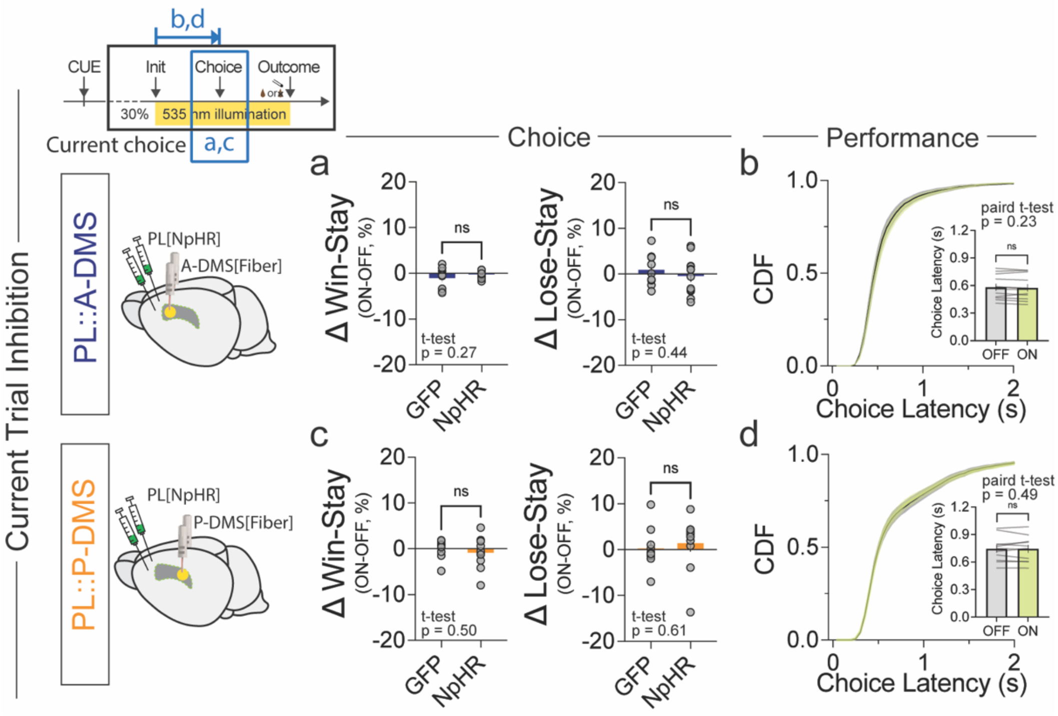


**Fig. S7. Optogenetic suppression of PL::A-DMS or PL::P-DMS activity during choice period does not affect current choice or motor performance.** a) Comparison of ΔWin-Stay (left) and ΔLose-Stay (right) between NpHR and GFP (control) groups, when light was delivered in the current choice epoch to A-DMS. b) Cumulative distribution of choice latencies and comparison of average choice latency (inset) from PL::A-DMS terminal illumination. c) Comparison of ΔWin-Stay (left) and ΔLose-Stay (right) between NpHR and GFP (control) groups, when light was delivered in the current choice epoch to P-DMS. d) Cumulative distribution of choice latencies and comparison of average choice latency (inset) from PL::P-DMS terminal illumination.

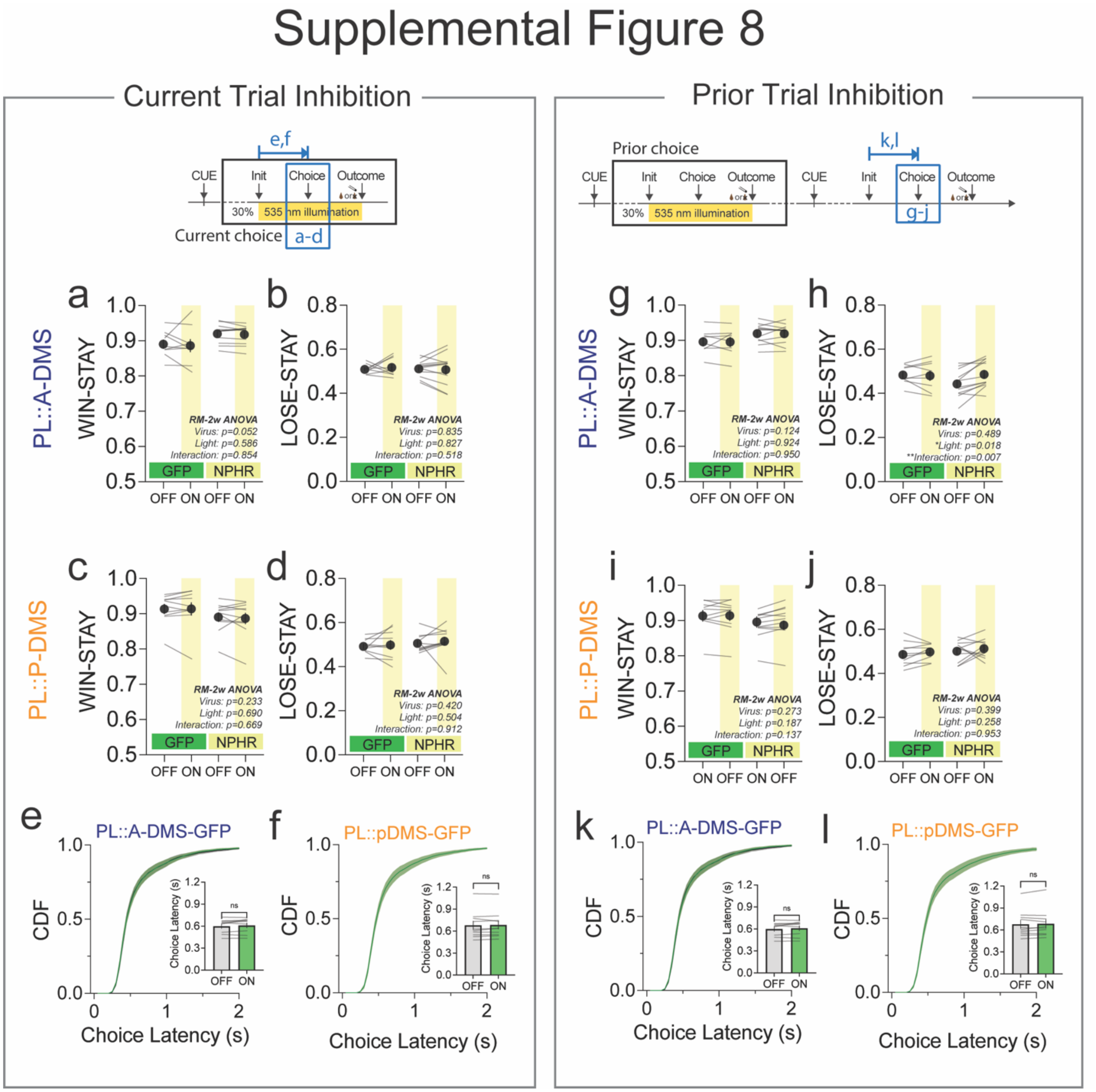


**Fig. S8. Effects of optogenetic suppression during choice period on behavior in current and subsequent trials.** a) Win-stay frequency in light ON/light OFF trials (paired comparison) for PL::A-DMS circuits infected with either GFP or NpHR. b) Lose-stay frequency in light ON/light OFF trials (paired comparison) for PL::A-DMS circuits infected with either GFP or NpHR. c) Win-stay frequency in light ON/light OFF trials (paired comparison) for PL::P-DMS circuits infected with either GFP or NpHR. d) Lose-stay frequency in light ON/light OFF trials (paired comparison) for PL::P-DMS circuits infected with either GFP or NpHR. e,f) Comparison of cumulative distribution of choice latency and average of choice latency (inset) in light ON versus OFF trials from (e) PL::A-DMS or (f) PL::P-DMS circuits injected with GFP. g-l) Identical layout as in (a-f) except all optogenetic manipulations were made on prior-trial.

### Supplemental Figure 9

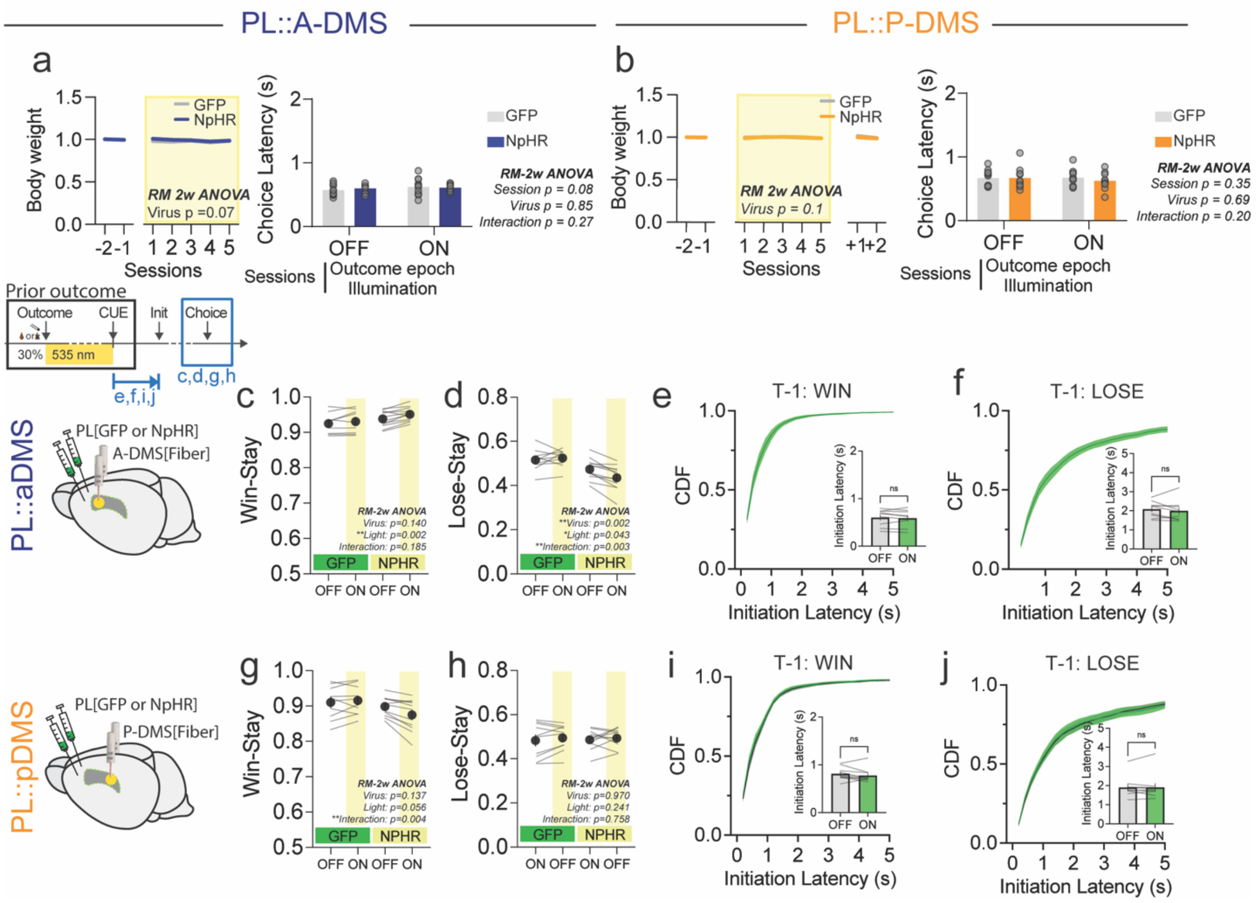


**Fig. S9. Effect of optogenetic suppression during outcome period on behavior.** a) Normalized body weight per session in sessions without and with random 30% outcome optogenetic inhibition of PL::A-DMS pathway (*left*). Comparison of choice latency between sessions with absence (OFF)/presence (ON) of outcome epoch illumination in PL::A-DMS from GFP or NpHR group (*right*). b) Normalized body weight per session in sessions without and with random 30% outcome optogenetic inhibition of PL::P-DMS pathway (*left*). Comparison of choice latency between sessions with absence (OFF)/presence (ON) of outcome epoch illumination in PL::P-DMS from GFP or NpHR group (*right*). c) Win-stay frequency in light ON/light OFF trials (paired comparison) for PL::A-DMS circuits infected with either GFP or NpHR. d) Lose-stay frequency in light ON/light OFF trials (paired comparison) for PL::A-DMS circuits infected with either GFP or NpHR. e,f) Cumulative distribution of initiation latencies and average initiation latency (inset) in light ON versus light OFF trials following (e) rewarded and (f) unrewarded outcomes for PL::A-DMS expressing GFP. g) Win-stay frequency in light ON/light OFF trials (paired comparison) for PL::P-DMS circuits infected with either GFP or NpHR. h) Lose-stay frequency in light ON/light OFF trials (paired comparison) for PL::P-DMS circuits infected with either GFP or NpHR. i,j) Cumulative distribution of initiation latencies and average initiation latency (inset) in light ON versus light OFF trials following (i) rewarded and (j unrewarded outcomes for PL::A-DMS expressing GFP.

### Supplemental Figure 10

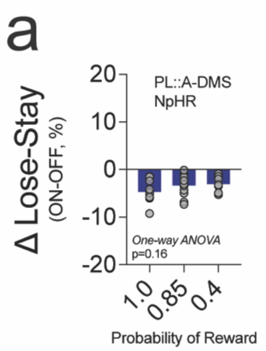


**Fig. S10. Similar behavioral consequences for PL::A-DMS inhibition across a range of reward probability paradigms.** a) ΔLose-Stay in different probability of reward.
